# The lncRNA *FENDRR* fine-tunes FOXF1 protein levels through a negative feedback loop governing human embryonic lung fibroblast-to-myofibroblast transition

**DOI:** 10.64898/2026.04.10.716765

**Authors:** Jean-François Laurendeau, Sophie Mockly, Gilberto Duran Bishop, Sophie Ehresmann, Elena Goretti, Virginie Calderon, Mohan Malleshaiah, Martin Sauvageau

## Abstract

Precise control of transcription factor dosage is critical for lung mesenchymal development. The forkhead box transcription factor FOXF1 is a dosage-sensitive regulator of pulmonary vascularization and fibroblast differentiation, with haploinsufficiency causing the lethal neonatal disorder alveolar capillary dysplasia with misalignment of pulmonary veins (ACDMPV). A syntenically conserved long noncoding RNA (lncRNA), *FENDRR*, is divergently transcribed ∼1.7 kb upstream of FOXF1. Notably, ACDMPV-associated chromosomal deletions disrupting *FENDRR* or a distal enhancer regulating both *FENDRR* and FOXF1 have been identified. Consistent with a role in lung development, selective deletion of *Fendrr* in mice causes neonatal lethality with lung growth defects, as well as alveolar and vascular abnormalities. Here, we define the conserved expression patterns, isoform diversity, and subcellular localization of *FENDRR* and FOXF1 across human and murine embryonic lung cell types. Functional perturbation reveals a bidirectional regulatory circuit in which FOXF1 promotes *FENDRR* transcription, while *FENDRR* limits FOXF1 protein abundance without affecting mRNA levels. Transcriptomic analyses demonstrate overlapping target gene networks and opposing effects on fibroblast-to-myofibroblast differentiation. These findings uncover a rheostat-like regulatory layer by which the lncRNA *FENDRR* fine-tunes FOXF1 protein dosage to influence lung fibroblast cell fate response and offers additional context for the significance of FOXF1-*FENDRR* dysregulation in ACDMPV.

## INTRODUCTION

The development of the respiratory system requires precise integration of transcription factors and regulatory elements to coordinate gene expression programs necessary for the proper alveolar branching and vascularization of lungs. Central to this process is FOXF1, a forkhead box transcription factor first expressed in the lateral plate mesenchyme surrounding the primitive gut and developing lung buds [1]. Within the developing lungs, FOXF1 is expressed in capillary endothelial cells, fibroblasts, and peribronchial smooth muscle cells and acts as a master regulator of mesenchymal differentiation, governing branching morphogenesis and vascular development [2, 3]. In endothelial cells, it governs VEGF signaling and receptor expression [3], while in fibroblasts, FOXF1 is critical for maintaining mesenchymal identity, regulating proliferation, and controlling myofibroblast activation [4–6].

Lung development is sensitive to FOXF1 dosage. In mice, homozygous loss causes mid-gestational lethality from mesoderm differentiation defects and complete vasculogenesis failure, while about half of heterozygous *Foxf1*^+/-^ mice die from pulmonary hemorrhage, lung hypoplasia, and defective alveolarization [7, 8]. Conditional endothelial overexpression of *Foxf1* similarly results in perinatal lethality accompanied by lung hypoplasia and a severely diminished capillary network, demonstrating that normal pulmonary development requires FOXF1 to be maintained within a certain quantitative window [9]. This dosage sensitivity is also observed in humans, where *FOXF1* haploinsufficiency and heterozygous mutations cause Alveolar Capillary Dysplasia with Misalignment of Pulmonary Veins (ACDMPV), a rare and neonatally lethal developmental lung disorder [10–12]. Affected newborns present with severe respiratory distress and intractable pulmonary hypertension within 48 hours of birth, with most dying within days to weeks despite supportive care. Approximately 80% exhibit concomitant cardiac, gastrointestinal, or genitourinary malformations [12]. Histologically, ACDMPV is defined by simplified lobular architecture, medial thickening of small pulmonary arteries, paucity of alveolar capillaries, abnormally positioned veins adjacent to pulmonary arteries within bronchovascular bundles, along with thickened alveolar septa, and, in some cases, increased interstitial fibroblasts and collagen deposition reflecting a fibrotic-type remodeling pattern [13–16].

The genetic basis of ACDMPV is characterized by heterogeneous spectrum of lesions centered on the 16q24.1 chromosomal locus. Approximately 40% of all diagnosed clinical cases are attributed to *de novo* heterozygous point mutations within the FOXF1 coding sequence [10]. Almost all remaining cases typically involve chromosomal deletions of variable size surrounding FOXF1. Notably, most large deletions that remove the FOXF1 gene also simultaneously delete the adjacent long noncoding RNA (lncRNA) *FENDRR*, transcribed in the opposite orientation only ∼1.7 Kb upstream [17]. Furthermore, nearly 40% of ACDMPV cases are caused by microdeletions that leave the FOXF1 coding sequence intact but selectively remove a distant lung-specific enhancer located ∼270 kb upstream [12, 18]. This regulatory element physically interacts with the intergenic promoter region via chromatin looping to coordinate the bidirectional transcription of both *FOXF1* and the lncRNA *FENDRR* [17]. The pathogenic relevance of this noncoding architecture is further evidenced by a subset of ACDMPV cases involving small, specific deletions that remove only the *FENDRR* gene or the shared intergenic promoter [17]. The allelic diversity observed in ACDMPV, supports the view that lung phenotype, histopathological severity, associated extrapulmonary anomalies, and, in some cases, atypical or attenuated clinical presentation is shaped not only by *FOXF1* haploinsufficiency itself, but also by the precise genomic context of each lesion. Such context may include the extent of enhancer disruption, involvement of neighboring *FENDRR* lncRNA, parental origin, mosaicism, or additional modifying variants, all of which could influence the severity and distribution of developmental defects in the lung.

The pathogenic impact of *FENDRR* deficiency is underscored by murine models where homozygous loss results in neonatal lethality due to lung lobular simplification evident from E14.5 (pseudoglandular stage), and later alveolar and vascular abnormalities reminiscent of ACDMPV [19, 20]. Adding to the functional importance of *FENDRR* in lung is the fact that it is downregulated in fibrotic lung fibroblasts from patients with idiopathic pulmonary fibrosis (IPF) [21]. Its ectopic expression in mouse lungs, using adenovirus, also reduces asbestos fibroblast-mediated fibrotic lesions and collagen deposition [22]. Meanwhile, FOXF1 mRNA and protein levels are upregulated in IPF-derived fibroblasts [23]. Several lncRNAs have been shown to regulate in *cis* neighboring protein coding genes [24, 25]. Despite the genomic proximity of *FOXF1* and *FENDRR*, the regulatory interplay between these two genes remains a subject of active investigation. Some models propose that *FENDRR* negatively regulates *FOXF1* mRNA levels in *cis* via repressive chromatin remodeling at its promoter, while genetic ablation studies report that *Fendrr* loss in mice does not significantly alter *Foxf1* mRNA levels in lungs, suggesting *FENDRR* acts through other mechanisms [19, 20]. Here, we contrast the conserved expression of both *FENDRR* and *FOXF1* across human and mouse embryonic lung cell types, their isoform diversity, and subcellular localization. We further show with antisense oligonucleotides (ASOs)-mediate depletions that human *FENDRR* negatively regulates FOXF1 protein levels, but not *FOXF1* mRNA. Supporting this, we demonstrate that both *FENDRR* and FOXF1 regulate the expression of common gene sets in opposing ways with concomitant impact on fibroblasts-to-myofibroblasts differentiation. Collectively, this study implicates *FENDRR* as a regulator of FOXF1 protein expression in human lung fibroblasts, revealing an unexpected layer of regulatory complexity between both genes.

## MATERIALS & METHODS

### Embryonic lung scRNA-seq analysis

Raw counts from mouse embryonic lung single-cell RNA sequencing (scRNA-seq) datasets (GEO GSE149563) were retrieved and processed as previously described [26]. For the human embryonic lung scRNA-seq dataset (ArrayExpress E-MTAB-11278) [27], the annotated expression matrix was directly retrieved and used for visualization. The Top 50 markers for each cell clusters are listed in Supplementary Tables S1 and S2 for human and mouse, respectively. The fibroblast population was further subdivided into proximal, mid, and distal subsets based on markers reported in the literature [28]. Time points corresponding to the pseudoglandular stage (E12 for mouse and W9 for human) were selected for analysis.

### Cell culture

WI-38 (CCL-75) and MLg (CCL-206) cell lines were obtained from the American Type Culture Collection (ATCC). Upon receipt, cells were initially grown, cryopreserved into multiple aliquots, and used from passage 4 through 12. Cell cultures were maintained in a humidified incubator at 37 °C, under 5% CO_2_, in DMEM (with 4.5 g/L glucose and L-glutamine, without sodium pyruvate; Wisent, 319-015-CS) supplemented with 10% heat-inactivated fetal bovine serum (FBS; Thermo Fisher Scientific, 12484028), 1x non-essential amino acids (Wisent, 321-011-EL), and 100 IU/ml of penicillin-streptomycin (Wisent, 450-201-EL).

### RNA isolation and RT-qPCR

Cell pellets (1-10x10^6^ cells) were first resuspended in 1ml of QIAzol (Qiagen, 79306) and RNA purified using the phenol–chloroform method [29]. For Illumina RNA-seq and direct RNA nanopore sequencing, RNA isolation was performed using Qiagen RNAeasy MiniKit columns (Qiagen, 74106) with modifications. Briefly, 200 µl of chloroform was added per 1 ml of Qiazol-sample mix, shook vigorously for 15 seconds and incubated for 3 min at room temperature. Samples were then centrifuged at 12,000 g for 15 min at 4°C, and the aqueous phase transferred to a clean RNAse-free tube. One volume of 75% ethanol was subsequently added, mixed, and transferred on an RNAeasy column. The remainder of the procedure was carried out as per the manufacturer’s protocol, including on-column DNase I (Qiagen, 79254) digestion. At the end, RNA was resuspended in RNase-free water.

Reverse transcription was performed with SuperScript IV (Thermo Fisher Scientific, 18091050) according to the manufacturer’s manual with oligo(dT) primers. Quantitative PCR (qPCR) analysis was carried out on a QuantStudio™ 5 Real-Time PCR System (Thermo Fisher Scientific) using PowerUp™ SYBR™ Green Master Mix (Thermo Fisher Scientific, A25741) with standard cycling parameters. A single reaction typically comprised 1 µM of primers with 10 ng of cDNA in a total volume of 10 µl. RNA levels were normalized using two housekeeping genes. Quantification was calculated using the ΔΔCt method [30]. See Supplementary Table S3 for qPCR primer sequences.

### Oxford Nanopore direct RNA long-read sequencing

RNA integrity was assessed using an Agilent 2100 Bioanalyzer with the RNA 6000 Pico Kit. Libraries were prepared from 2 µg of total RNA using the Direct RNA Sequencing Kit (Oxford Nanopore Technologies, SQK-RNA004), including cDNA synthesis with Induro Reverse Transcriptase (NEB, M0681). Each library was sequenced on an PromethION™ RNA flow cell (FLO-PRO004RA), yielding 15.07 M reads (N50 = 2.32 kb) for MLg and 12.03 M reads (N50 = 2.04 kb) for WI-38. Raw POD5 data were base called with Dorado v0.9.5 (Oxford Nanopore Technologies) using the super-accuracy model (rna004_130bps_sup@v5.1.0), and FASTQ files were aligned to the human GRCh38 (release 46) or mouse GRCm39 (release M33) primary assemblies from GENCODE using Minimap2 v2.28 [31] with the splice-aware parameters ‘--ax splice –uf’. Isoform identification was performed with StringTie3 v3.0.0 [32] in long-read mode ‘-L’ without reference annotation, and *FENDRR* isoforms were curated manually. The resulting transcript annotation was then used as a reference for isoform quantification with StringTie3 in expression estimation mode ‘-e’.

### Single-molecule fluorescence *in situ* hybridization (smFISH)

Probes (20-mers) were designed using the Stellaris Probe Designer tool v4.2. Selection criteria included 30%-60% GC content and spacing length of at least 2 nucleotides between oligos. Oligos were further curated using BLAST to minimize risk of off-targets. Selected probes (Supplementary Table S4) were enzymatically synthesized and Deep Red (647)-labelled by DNA Script using their SYNTAX™ system technology. Probes were resuspended in DNase/RNase-free water at a concentration of 5 µM. Probe sets were prepared at 20 ng/ul in nuclease-free water and aliquots stored at -20°C protected from light. Single-molecule FISH was performed as previously described [33] with modifications. Round 18 mm coverslips were coated with bovine type I collagen (Stemcell Technologie, 07001) in six-well plates. WI-38 or MLg cells were seeded at 2.5x10^5^ cells per well and grown for 48h. Cells were subsequently washed with PBS (Wisent, 311-425-CS) and then fixed for 10 min with fresh 4% formaldehyde in PBS. Next, cells were washed twice with PBS and kept at -20°C in 70% ethanol until hybridisation. On the day of hybridisation, cells were washed twice with wash buffer: 10% deionized formamide (Millipore Sigma, F9037-100ML) and 2x SSC (300 mM NaCl, 30 mM sodium citrate). In parallel, probes were prepared by mixing 20 ng of each probe sets with 40 µg of sheared salmon sperm DNA (Thermo Fisher Scentific, AM9680) and Yeast tRNA (Thermo Fisher Scentific, 15401011). Probe mixes were then dried with vacuum centrifugation and reconstituted in 20 µl of hybridization buffer: fresh 10% deionized formamide, 10% dextran sulfate (Bio Basic, DB0160-10), and 2x SSC. Next, reconstituted probes were heated for 3 min at 95°C, after which 2 µg of BSA (Wisent, 800-095-EG) in aqueous solution as well as 40 pmol of 65°C-preheated Ribonucleoside Vanadyl Complex (New England Biolabs, S1402S) were added. Coverslips were then incubated with probe mixes in a dry hybridisation chamber for 3h at 37°C in the dark. After hybridization, samples were rinsed once with wash buffer for 30 min at 37°C in the dark, followed by two PBS washes. Finally, coverslips were dipped in 100% ethanol, air dried for 30 sec until cells appeared white, and mounted on slides with ProLong Gold Antifade Mountant containing DAPI (Thermo Fisher Scientific, P36931). The mounting medium was allowed to dry overnight, after which coverslips were immediately sealed with nail polish and images acquired.

Fluorescence confocal microscopy was performed on a Leica Stellaris confocal microscope with TauSense lifetime-based imaging technology and a 63x/1.4 oil immersion objective (Leica, 506350). Samples were illuminated using a 405 nm diode laser for DAPI and a 653 nm wavelength selected from a tunable white light laser for smFISH probes. Gain and offset settings were adjusted to avoid pixel saturation. Tau gating was applied using a lifetime window of 0.3–8.0 ns and images were subjected to line accumulation twice, with an average pixel dwell time of 0.5125 μs. Image processing was done using Fiji (ImageJ2 v2.14.0) [34]. Confocal images represent maximum intensity projections of z-stacks. For visualization purposes, a Gaussian blur (σ = 2 pixels) was applied uniformly to all images, and brightness and contrast were adjusted

### Subcellular fractionation

Ten million cells were resuspended in lysis buffer (10 mM Tris-HCl pH 7.9, 100 mM KCl, 5 mM MgCl_2_, 1 mM DTT) and incubated on a rotating platform at 4°C for 5 min to ensure thorough mixing. The cell lysate was then carefully layered on top of nuclear centrifugation buffer (50% glycerol, 20 mM Tris-HCl pH 7.4, 75 mM KCl, 5 mM MgCl_2_, 1 mM DTT) and centrifuged at 2,000 g at 4°C for 4 min. The cytoplasmic supernatant was transferred to a new tube, and the nuclear pellet was washed with lysis buffer and centrifuged again. Thereafter, the supernatant was discarded, and the nuclei pellet resuspended in glycerol/urea buffer (25% glycerol, 20 mM Tris-HCl pH 7.4, 187.5 mM KCl, 7.5 mM MgCl_2_, 0,5 M urea, 10% IGEPAL CA-630, 1 mM DTT). Nuclei were allowed to lyse for 5 min on ice, followed by centrifugation at 2,000 g for 4 min at 4 °C. The nucleoplasmic supernatant was then transferred to a new tube, and the chromatin pellet was resuspended in lysis buffer. To isolate RNA, lysates were mixed 1:1 with QIAzol reagent, and RNA purification was performed as described above. RNA from each fraction was resuspended in an equal volume of RNase-free water. Equal volumes from each fraction were also used for downstream RT-qPCR analysis, and individual RNA targets were presented as their distribution across subcellular fractions.

### Sequence Alignments

FASTA format of human *FENDRR1* and *FENDRR2* as well as mouse *Fendrr* sequences, derived from nanopore direct RNA sequencing data of WI-38 and MLg cells, were used as input. Pairwise sequence alignments were performed using Clustal Omega (v1.2.4) via the EMBL-EBI web server (https://www.ebi.ac.uk/Tools/msa/clustalo/). Nucleotide sequences were submitted with the following parameters: mBed-like clustering with 5 combined guide tree iteration, max HMM iterations 5, distance calculation via k-tuple similarity with k=6, gap penalties of 4.0 for open and 4.0 for extension. Alignments were exported as aligned FASTA and CLUSTAL formats for downstream analysis of sequence conservation.

### Antisense oligonucleotide (ASO) knockdown

Two non-targeting control gapmer ASOs, two gapmer ASOs targeting *FENDRR*, and two gapmer ASOs targeting *FOXF1* were used in the current study (see Supplementary Table S5 for sequences). All gapmer ASOs (custom design, IDT) are 20 nucleotides long, 5′-labeled with 6-carboxyfluorescein (6-FAM), fully modified with phosphorothioate linkages and 2′-O-methyl modifications in a 5-10-5 design. Gapmer ASOs were transfected with Lipofectamine RNAiMAX (Thermo Fisher Scientific, 13778150) according to the manufacturer’s guidelines at a final concentration of 50 nM. Unless otherwise indicated, cells were typically transfected in a forward manner, with cells being seeded at 41.7x10^3^ cells/cm^2^ the day before and collected 48h following transfection. Afterward, cells were washed with PBS, detached with Trypsin (Wisent Bioproducts, 325-044-EL), resuspended in complete media, and pelleted by centrifugation at 400 g for 5 min. Cells were then resuspended in PBS with 1% FBS and sorted with fluorescence activated cell sorting (FACS). Sorted cells were subsequently pelleted by centrifugation at 400 g for 5 min, and either resuspended in Qiazol or in RIPA buffer (Bio Basic, RB4475) with protease inhibitor (Sigma-Aldrich, 11697498001) for RNA-seq or western blotting, respectively.

### RNA sequencing

RNA integrity was first assessed on a 2100 bioanalyzer (Aglient Technologies) using the RNA 6000 Pico Kit. Libraries were then prepared from 172 ng total RNA using polyA enrichment with the NEBNext Poly(A) mRNA Magnetic Isolation Module (New England Biolabs) followed by preparation with the RNA Hyperprep Kit (KAPA). Library size distribution was assessed on a 2100 bioanalyzer (Agilent Technologies) using a High Sensitivity DNA Kit and library quantification performed by qPCR. Equimolar libraries were sequenced in paired-end reads (PE100) on a Novaseq 6000 system (Illumina) with an S4 flowcell at a coverage of ∼30M fragments per library. Read mapping was done with STAR v2.7.9a aligner [35] with the twopassMode Basic mode and the Gencode V42 annotation. Quantification was performed with featureCounts v1.6.4 [36], using the same annotation. DESeq2 v1.44.0 [37] was used to normalize read counts and identify differentially expressed genes (DEGs). Genes were considered differentially expressed if their adjusted p-value (Padj) was below 0.05 and if z-scores of their normalized counts showed a consistent direction of change across both target ASOs, thereby excluding non-specific effects of single ASO. DEGs are listed in Supplementary Table S6 and Supplementary Table S7. GO terms were analyzed with msigdbr v10.0.2 and the Molecular Signatures Database (MSigDB) gene sets [38]. The RNA-seq data are deposited in the Gene Expression Omnibus (GEO).

### Triplex forming oligonucleotide analysis

Triplex-Forming Oligos (TFO) and Triplex-Target Sites (TTS) were identified using the *Triplex Domain Finder* (rgt-TDF) algorithm [39, 40]. Briefly, FASTA sequences of human *FENDRR* isoforms identified by Nanopore direct RNA sequencing were extracted. BED files for promoters of differentially expressed genes upon *FENDRR* knockdown, along with background promoters from all expressed genes in non-targeting and *FENDRR* ASO conditions, were generated using R packages rtracklayer and GenomicRanges based with the hg38 human genome assembly and GENCODE v42 annotation. Promoter sequences were defined as sequences 2 kb upstream of the TSS to 500 bp downstream of the TSS. The rgt-TDF promotertest was then run with base parameters and significance threshold –a 0.1. TFO localisation was then realigned to the genome using R package GenomicRanges and visualized using Gviz.

### Western blotting

Cells were lysed in RIPA buffer (Bio Basic, RB4475) with protease inhibitor (Sigma-Aldrich, 11697498001). Lysates were separated on 12.5% SDS-PAGE gels and transferred to polyvinylidene difluoride (PVDF) membranes using Tris-glycine-ethanol transfer buffer (25 mM Tris, 192 mM glycine, 20% ethanol). Membranes were blocked for 1h in 5% skim milk diluted in TBST (198 mM Tris, 1506 mM NaCl, 0.1% Tween-20), and all antibody incubations were performed in the same buffer, interspersed with four TBST washes. Blots were incubated overnight at 4 °C with fresh anti-FOXF1 antibody (1 µg/mL; Bio-Techne, AF4798), followed by 1h incubation with HRP-conjugated donkey anti-goat secondary antibody (1:7,000; Jackson ImmunoResearch, 705-035-003). Signal was developed using enhanced chemiluminescence (ECL) and imaged using the Chemi Hi Sensitivity mode on a Bio-Rad ChemiDoc MP Imaging System. Membranes were subsequently re-probed for loading control using anti-α-Tubulin primary antibody (1:7,000; Proteintech, 11224-1-AP) and HRP-conjugated goat anti-rabbit secondary antibody (1:10,000; Bio-Rad, 170-6515).

### FOXF1 ChIP-seq and FOXF1 motif analysis

Raw sequencing reads from FOXF1 ChIP-seq embryonic mouse lung dataset (GEO GSE77951) [9] were retrieved and trimmed using Trimmomatic v0.39 [41]. Reads were aligned to the GRCm39 reference genome using Bowtie2 v2.5.3 [42], with an average of 75% uniquely mapped reads. MACS3 v3.0.1 [43] was used to call significant peaks in narrowPeak mode (adjusted p-value < 0.05). Consensus peaks across replicates were defined using BEDtools v2.31.1 [44], and peak quantification was performed with featureCounts v2.0.6 using the Gencode VM37 annotation. Read counts were normalized with DESeq2 v1.41.1, and peak annotation was carried out using HOMER v4.11.1 [45] with the nearest feature. DEGs from our RNA-seq experiments were converted to mouse gene symbols using BioMart, retaining only one-to-one orthologs, and were intersected with the FOXF1 ChIP-seq dataset (See Supplementary Tables S8 and S9). Similarly, FOXF1 binding motifs, as determined by JASPAR, were identified in human gene promoters, defined as -2 Kb to +2 Kb (GENCODE V42 annotation, hg38) and intersected with *FENDRR* and *FOXF1* DEGs and the FOXF1 ChIP-seq dataset.

### SHH and TGFβ1 treatment

Cells were seeded at a density of 72.9 × 10^3^ cells/cm^2^ and allowed to adhere and grow for 24h. Cells were then starved for an additional 24h in complete medium containing 1% FBS, and starvation conditions were maintained for the remainder of the experiment. One control well was left untreated throughout. For Sonic Hedgehog (SHH) pathway induction, cells were either pre-treated with 135 nM of SHH antagonist SANT1 (Sigma-Aldrich, 559303) for 20 min or left untreated, followed by supplementation with recombinant SHH (Bio-Techne, 1845-SH) at a final concentration of 10 nM. For TGFβ pathway induction, cells were directly treated with recombinant TGFβ1 (BioLegend, 781802) at a final concentration of 10 ng/mL. After 48h of SHH or TGFβ stimulation, cells were harvested in QIAzol for downstream RT-qPCR analysis.

### Wound healing assay

The day prior to wound making, Incucyte Imagelock 96-well plates (Sartorius, 4379) were coated with bovine type I collagen. Cells were reverse transfected with ASOs at a final concentration of 50 nM and seeded at 80x10^3^ cells/well. Cells formed a 100% confluent monolayer 24h later. At this moment, transversal wounds were generated in each well using the Incucyte 96-Well Woundmaker Tool (Sartorius, 4474) following the manufacturer’s guidelines (publication no. 8000-0597-B00). Images were acquired every hour over a 40h period using the IncuCyte platform in scratch wound scan mode at 10x magnification, capturing phase contrast. Analysis was performed with the IncuCyte Scratch Wound Analysis software using the built-in phase analysis to detect and segment cells. The wound confluence was calculated as the percentage of the wound region area occupied by cell.

### Immunofluorescence staining and imaging

Cells were seeded on Ibidi µ-Slide 8 Well with ibiTreat (ibidi, 80826) at 73x10^3^ cells/well. For depletion assays, cells were reverse transfected with ASOs at a final concentration of 50 nM prior seeding. TGFβ pathway induction was performed as described above. Fibroblast-to-myofibroblast transition was assessed either 24h or 48h after addition of TGFβ1 for depletion or overexpression assays, respectively. At endpoint, cells were washed twice with PBS, fixed with 4% formaldehyde in PBS for 10 min, and washed an additional three times with PBS. Cells were then stored at 4°C in PBS until probing. On the day of immunostaining, cells were first permeabilized for 10 min with 0.3% Triton X-100 in PBS. Cells were subsequently washed three times in PBS, followed by 1h incubation with blocking buffer (1% BSA, 0.3% Triton X-100, 1X PBS). Afterward, cells were incubated 2h at room temperature with anti-α-Smooth Muscle Actin (αSMA 1:1000 dilution; Sigma-Aldrich, A5228), followed by three PBS washes and an additional 1h incubation with AlexaFluor647-conjugated anti-mouse secondary antibody (Thermo Fisher Scientific, a21235). Finally, cells were incubated 5 min in PBS with 1 µM DAPI, preceded and followed by three PBS washes. Cells were kept in PBS at 4°C until imaging.

Fluorescence confocal microscopy was performed on a Leica Stellaris confocal microscope with a 20x/0.75 dry objective (Leica, 506517). Samples were illuminated using a 405 nm diode laser for DAPI and a 653 nm wavelength selected from a tunable white light laser for detection of AlexaFluor647-conjugated antibodies. Gain and offset settings were adjusted to avoid pixel saturation. Images were subjected to 4 times line averaging, with an average pixel dwell time of 2.8 μs. Multiple images per well were acquired. Image processing was done using Fiji. Confocal images represent maximum intensity projections of z-stacks. A pixel intensity cutoff was set to substract background and generate regions of interest (ROI). Integrated intensity of the αSMA signal was then measured across samples. Brightness and contrast were adjusted for visualization purposes only.

### Lentiviral expression

Two isoforms of *FENDRR*, identified by long read nanopore sequencing, were selected for ectopic overexpression (see Supplementary Table S10), with each corresponding to the most expressed isoform of the 3’ proximal and distal end. Constructs were generated by GenScript, inserting each *FENDRR* isoform sequence into a pLenti-CMV-GFP-Hygro backbone. Lentiviral particles were produced and used to infect passage five WI-38 cells at a multiplicity of infection (MOI) of 1. Following infection, GFP^+^ cells were isolated by FACS to obtain an exclusive population of *FENDRR*-overexpressing cells. These cells were then expanded, cryopreserved in multiple aliquots, and used for experiments between passages 8 and 12 post-transduction.

### Statistics and reproducibility

Statistical analyses were performed using the R software. One-way ANOVA and Student’s t-test were used to assess statistical significance, with a significance threshold set at *p* < 0.05. Two-way ANOVA was used for time-course experiments (*i*.*e*. proliferation and migration assays). When applicable, Dunnett’s post-hoc test was performed to compare each treatment group to the control. Data are presented as mean ± standard deviation (SD). All experiments were performed with three independent biological replicates or more.

## RESULTS

### *FENDRR* and *FOXF1* are co-expressed in specific lung embryonic subtypes

In mice, the lncRNA *Fendrr* was previously found to be co-expressed with its neighboring protein coding gene *Foxf1* in the caudal end of the lateral plate mesoderm of mid gestation embryos [46], and in the mesoderm of embryonic lungs [19]. Since genetic deletion of *Fendrr* results in lung developmental defects observable at the pseudoglandular stage [19, 20], we reanalyzed publicly available human and mouse embryonic lung single-cell RNA-seq datasets, focusing on week 9 of gestation for human and E12 for mouse to get a more precise profile of which specific lung cell types express *FENDRR* during development [26, 27]. After reproducing the original cell clustering, we further subdivided the fibroblast population into proximal, mid, and distal subsets based on literature-reported markers [28]. In human scRNA-seq datasets, *FENDRR* is expressed in fibroblasts, myofibroblasts, smooth muscle cells (SMC), as well as vascular endothelial, and mesothelial cells (Fig.1A,C). Similarly in mice, *Fendrr* is expressed in fibroblasts, secondary crest myofibroblasts, smooth muscle and mesothelial cells, as well as proliferating and capillary endothelial cells (Fig.1B,C). *FOXF1* displays a similar expression pattern as *FENDRR* in both human and mouse (Fig.1A-C). Within the fibroblast subpopulations, *FENDRR* and *FOXF1* expression appears to be higher in proximal cells and decreasing towards distal positions (Fig.1A-C), which would be consistent with proximal fibroblasts being closer to epithelial cells which secrete Sonic Hedgehog (SHH), a known activation of *FOXF1* expression [47, 48]. While being expressed in the same cell types, *FOXF1* expression was detected in a higher number of cells within most subtypes. Accordingly, when looking at co-expression, the majority of *FENDRR*^+^ cells express *FOXF1* as well (Fig.1D). This is particularly true for proximal fibroblasts and other non-fibroblast cells. In contrast, detection of both *FENDRR* and *FOXF1* in distal and mid fibroblast appear less frequent.

**Figure 1.**
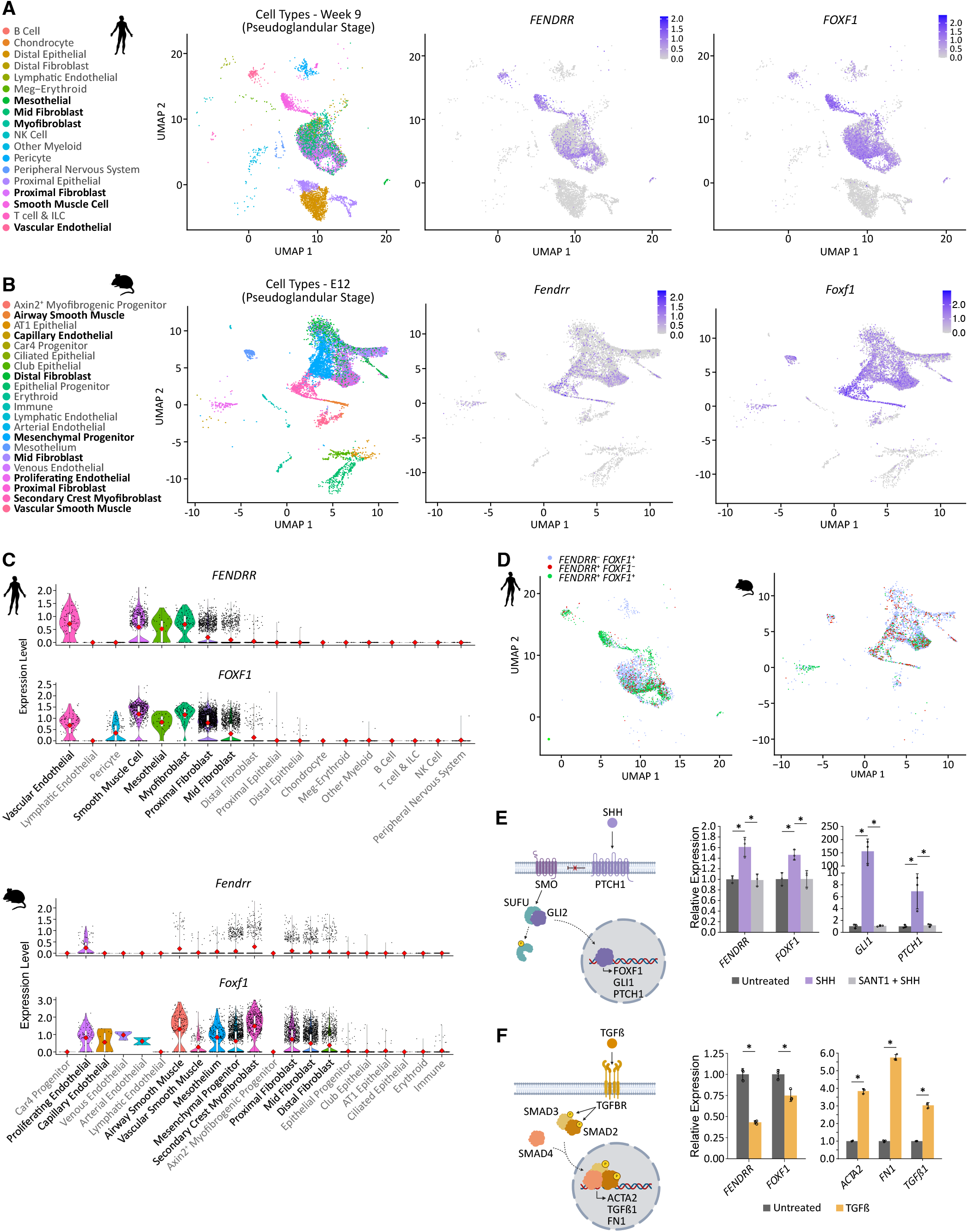
*FENDRR* and *FOXF1* are co-expressed in human and mouse embryonic lung mesenchymal subpopulations. (**A**,**B**) scRNA-seq UMAPs showing RNA expression levels of *FENDRR* and *FOXF1* in specific mesenchymal cell lineages of human week 9 (**A**) and mouse E12 (**B**) embryonic lungs at the pseudoglandular stage. (**C**) Violin plots showing RNA expression levels of human (top) and mouse (bottom) *FENDRR* and *FOXF1* across embryonic lung cell types. Each dot represents a expression levels in a cell. (**D**) UMAPs of human week 9 (left) and mouse E12 (right) embryonic lung cell mesenchymal subsets co-expressing *FENDRR* and *FOXF1*. (**E**) RT-qPCR analysis of *FENDRR* and *FOXF1* expression 48h after Sonic Hedgehog (SHH) stimulation, with or without prior treatment with the SANT1 inhibitor, in human WI-38 embryonic lung fibroblasts. *GLI1* and *PTCH1* expression confirms effective activation (SHH) and inhibition (+SANT1) of SHH signaling. Expression levels were normalized to the geometric mean of the housekeeping genes *TBP* and *GAPDH*, and fold changes were calculated relative to untreated cells. Bars represent the mean of untreated (dark grey), 10nM SHH-treated (purple), or 135 nM SANT1-pretreated (SHH antagonist; light grey) cells. Dots indicate individual biological replicates (n=3), and error bars denote standard deviation (SD). Statistical significance was determined by one-way ANOVA followed by Dunnett’s post hoc test (* *p*-val < 0.05). (**F**) RT-qPCR analysis of *FENDRR* and *FOXF1* expression 48h after TGFβ1 induction in WI-38. Expression of *ACTA2, FN1*, and *TGFB1* confirms activation of TGFβ1 signaling. Expression values were calculated, plotted, and statistically analyzed as in E. Bars represent the mean of untreated (dark grey) and TGFβ1-treated (yellow) cells.

Since *FENDRR* and *FOXF1* appear to be largely co-expressed in proximal fibroblasts, endothelial, and smooth muscle cells, we examined whether *FENDRR* and *FOXF1* were co-regulated by upstream regulatory pathways, such as SHH and TGFβ1, which are known to regulate these cell types during lung development [48–53]. To assess co-regulation, we decided to induce the SHH and TGFβ pathways in WI-38 embryonic lung fibroblasts, which are derived from a 3-months old fetus (lung pseudoglandular stage). Treatment of WI-38 cells with SHH induced *FENDRR* expression level alongside *FOXF1*. As expected, this induction was reverted when the antagonist of SHH signaling, SANT1, was added to cells (Fig.1E). Conversely, exposing cells to TGFβ1 inhibited both *FENDRR* and *FOXF1* expression at the RNA level (Fig.1). Together, these data shows that *FENDRR* and *FOXF1* are co-expressed in the same lung mesenchymal and mesodermal subpopulations in both human and mouse and are transcriptionally co-regulated at least in human embryonic lung fibroblasts by SHH and TGFβ1.

### Human and Mouse *FENDRR* show isoform and subcellular localization diversity

*FENDRR* and *FOXF1* loci are positionally conserved in both species, with their transcription start sites (TSSs) separated by approximately 1.7 kb on chromosome 16 and 1.2 kb on chromosome 8 in human and mouse, respectively (Fig.2A,B). As indicated by phylogenetic codon substitution frequencies analysis using PhyloCSF) [54], both human and mouse *FENDRR* lack coding potential, and are enriched for marks of active transcription, namely H3K4me3 and H3K27ac, at their promoters in human lung fibroblasts and mouse lungs (Fig.2A,B). Current GENCODE annotations contain several *FENDRR* isoforms. To determine which are expressed in human and mouse lung fibroblasts, we utilized the human embryonic lung fibroblast WI-38 and mouse newborn lung fibroblast MLg cell lines, which express *FENDRR* and *FOXF1* at appreciable levels. Using these, we performed direct RNA long read sequencing with Oxford Nanopore technology to resolve full-length isoforms. All identified transcripts for both human and mouse initiated within the first annotated exon in accordance with CAGE-seq peaks, with no evidence of alternative transcription start sites (Fig.2A,B). Human *FENDRR* showed greater isoform diversity compared to mouse, especially at the 3’ end, where alternative splicing generates two groups of *FENDRR* transcripts, one ending at a proximal 3’ end and another at a more distal 3’ end, which compose the majority of *FENDRR* sequences (Fig. 2A). Among these, only 7/17 GENCODE annotated isoforms were detected in WI-38 cells (Fig.2A, dark red), whereas we identified 7 novel isoforms (Fig.2A, light red). Transcript quantification revealed that in human *FENDRR* transcripts with the proximal 3’ end are usually more expressed than those with the distal 3’ end. Two of these transcripts, one with a proximal 3’ end and one with a distal 3’ end were cloned in lentiviral vectors for subsequent functional experiments (Fig.2C). In mice, none of the GENCODE annotated isoforms were detected in MLg cells (Fig.2B, grey). Instead, we found 2 distinct transcripts that differed only at their 3’ end (Fig.2B, light red), with the longest isoform being dominantly expressed (Fig.2D).

**Figure 2.**
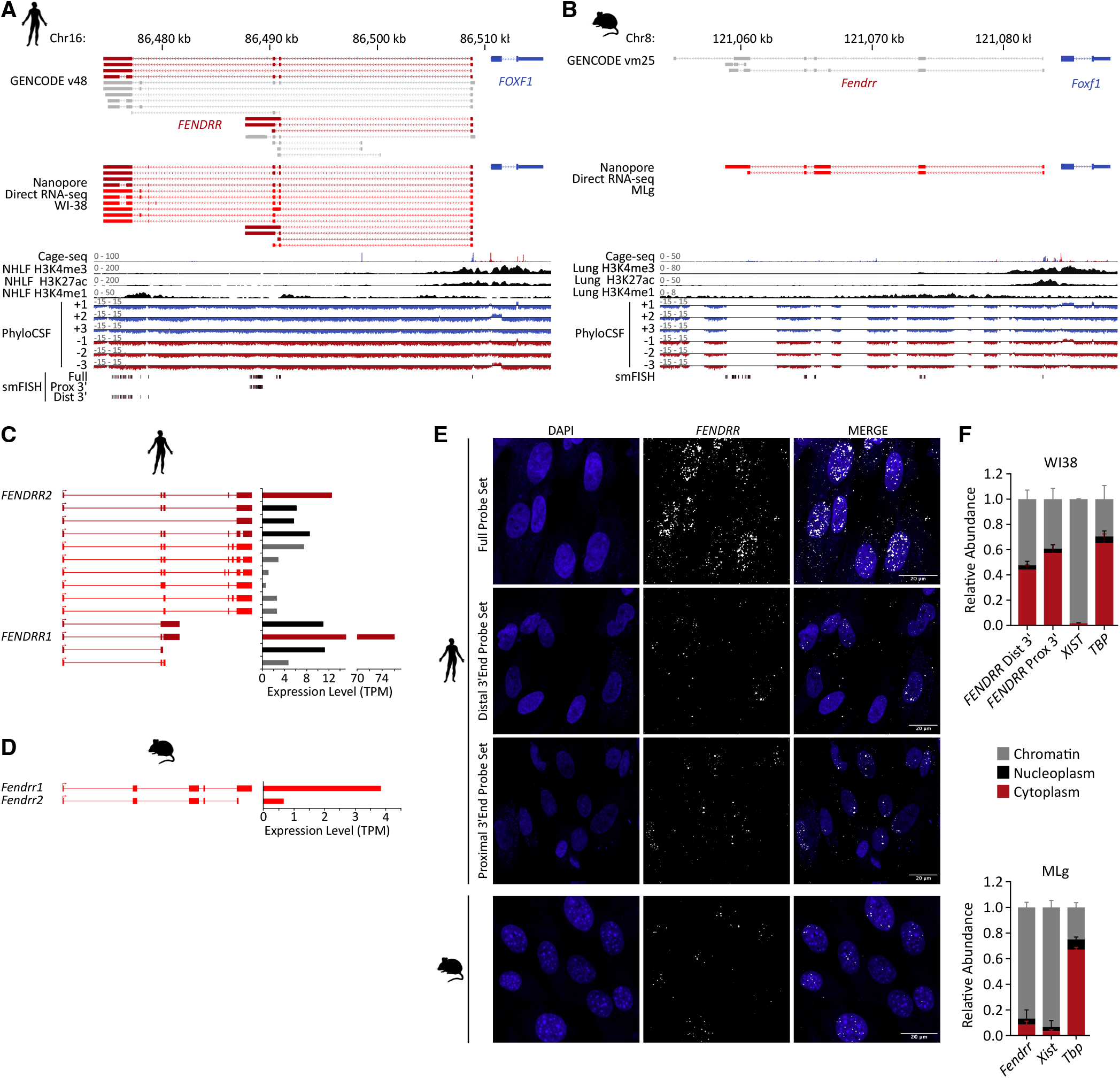
Human and mouse *FENDRR* isoform expression, subcellular localization, and conservation. (**A**,**B**) Genomic loci of *FENDRR* and *FOXF1* in human (**A**) and mouse (**B**). Comparison of GENCODE human v48 (hg38) or mouse vm25 (mm10) transcript annotations and direct RNA nanopore sequencing performed in WI-38 and MLg cells identifies known (dark red) isoforms, newly identified isoforms (light red), and undetected isoforms (ligh grey). FANTOM5 Cage-seq peaks indicate transcription start sites (TSS), together with H3K4me3, H3K27ac, and H3K4me1 peaks from normal human lung fibroblasts (NHLF; **A**) or mouse lungs (**B**). PhyloCSF tracks across the six reading frames indicate lack of coding potential for human and mouse *FENDRR*. The positions of the single-molecule RNA FISH (smFISH) probe sets used in this study are indicated for reference. (**C**) Quantification of human *FENDRR* isoform expression in WI-38 embryonic lung fibroblasts. *FENDRR* isoforms identified by Nanopore direct RNA sequencing were quantified in transcripts per million (TPM) using StringTie3. The two isoforms cloned into lentiviral expression vectors are indicated as *FENDRR1* and *FENDRR2*. (**D**) Quantification of mouse *Fendrr* isoform expression in MLg neonatal lung fibroblasts. *Fendrr* isoforms identified by Nanopore direct RNA sequencing were quantified in transcripts per million (TPM) using StringTie3. (**E**) smFISH detection of human and mouse *FENDRR* in WI-38 and MLg cells, respectively. For human *FENDRR*, different probe combinations were used to detect all *FENDRR* transcripts (top) or only transcripts containing the distal 3’ end (middle) or proximal 3’ end (bottom). (**F**) Relative RNA abundance of human and mouse *FENDRR* in cytoplasmic, nucleoplasmic, and chromatin subcellular fractions, as determined by RT-qPCR in WI-38 cells (top) and MLg (bottom) cells. *XIST* and *TBP* were used as chromatin and cytoplasmic control genes, respectively.

Next, we performed single-molecule RNA fluorescence *in situ* hybridization (smFISH) using different probe sets to target human *FENDRR* transcripts. An initial full probe set was used to detect signal from all *FENDRR* isoforms (Fig. 2A,E). Probes were then split to create sets that would recognize either *FENDRR* isoforms with the proximal 3’ end or the distal 3’ end (Fig.2A). All showed that human *FENDRR* localizes both in the nucleus and the cytoplasm in WI-38 cells (Fig. 2E). We further confirmed these findings by performing subcellular fractionation followed by RT-qPCR with different primer pairs. Similar to smFISH, fractionation showed that transcripts with proximal or distal 3’ end were distributed among the chromatin and the cytoplasmic fractions (Fig. 2F, top). In contrast, smFISH and subcellular fractionation with RT-qPCR revealed that mouse *Fendrr* was predominantly localized in the nucleus and chromatin fraction, respectively (Fig. 2B,E,F). These findings indicate that human *FENDRR* acquired increased isoform diversity and exhibits distinct localization patterns compared to mouse.

### *FENDRR* negatively regulates FOXF1 protein expression

Because of its association with ACDMPV, we were interested in investigating further the role of *FENDRR* in human lung fibroblasts and clarify its potential interplay with FOXF1. First, we used gapmer antisense oligonucleotides (ASOs) to knockdown either *FENDRR* or *FOXF1* in WI-38 cells. For each gene, two ASOs were designed to target all isoforms while being located further downstream of the first exon to minimize the risk of premature transcription termination by RNA polymerase II and potential act of transcription effects [19, 55, 56]. ASO-mediated knockdown (KD) of *FENDRR* or *FOXF1* resulted in a ∼70% and ∼81% decrease in RNA expression levels assessed by RT-qPCR, respectively (Fig.3A). *FOXF1* KD was further validated by western blot, showing a near-complete loss of the protein (Supplementary Fig.S1A). In accordance with *Fendrr* knockout mice [19], ASO-mediated *FENDRR* depletion did not significantly alter *FOXF1* expression at the RNA level (Fig.3A, left). However, we did observe a significant ∼31% increase in FOXF1 protein levels by western blot in *FENDRR* KD WI-38 cells compared to non-targeting ASO controls (Fig.3B). Conversely, a significant ∼25% decrease of *FENDRR* expression was observed upon *FOXF1* depletion (Fig.3A, right), in accordance with reports that FOXF1 mutations leads to decreased *FENDRR* levels in ACDMPV patient samples [57].

**Figure 3.**
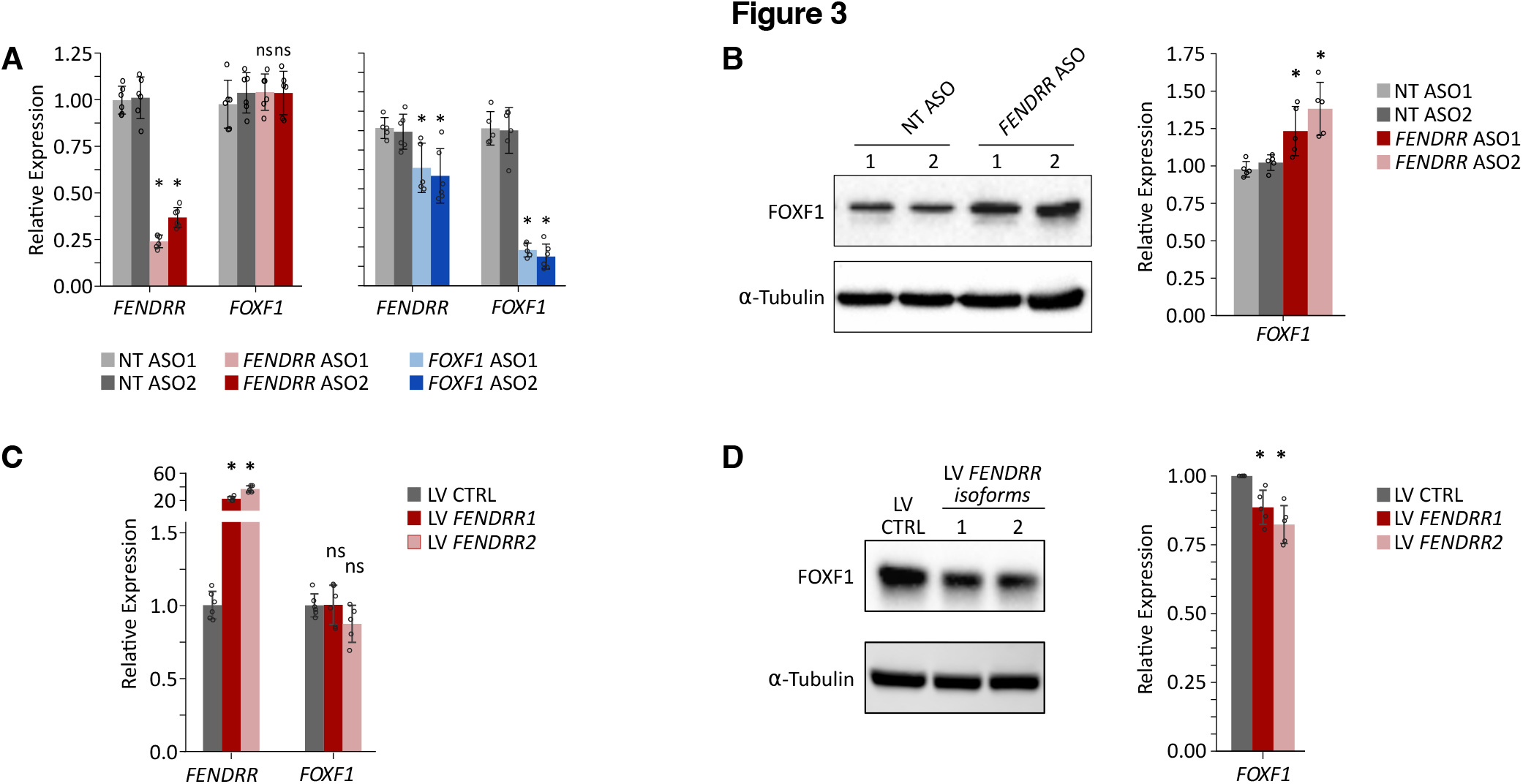
*FENDRR* negatively regulates FOXF1 protein, but not RNA levels. (**A**) RT-qPCR analysis of *FENDRR* and *FOXF1* expression 48h after ASO-mediated knockdown of *FENDRR* (left; red) or *FOXF1* (right; blue) in WI-38 cells. Fold changes were calculated relative to both non-targeting control antisense oligonucleotides (NT ASO1 and NT ASO2). Each ASO pair was tested in six biological replicates (dots, n=6). Bars represent the mean of individual ASOs, and error bars indicate standard deviation (SD). Statistical significance was determined by one-way ANOVA followed by Dunnett’s post hoc test (* *p*-val < 0.05; ns, not significant). (**B**) Representative western blot showing FOXF1 protein levels (left) and corresponding quantification across five biological replicates (right). FOXF1 protein abundance was normalized to the housekeeping protein α-Tubulin. Fold changes were calculated and plotted as in A. Statistical significance was determined by one-way ANOVA followed by Dunnett’s post hoc test (* *p*-val < 0.05). (**C**,**D**) *FENDRR* and *FOXF1* RNA (**C**) and FOXF1 protein (**D**) expression levels measured by RT-qPCR and western blot, respectively, after lentiviral overexpression of *FENDRR* isoforms in WI-38 cells.

To further test this with an orthogonal overexpression approach, we cloned in lentiviral vectors the most highly expressed *FENDRR* isoform containing a proximal 3’ end, which we named *FENDRR1*, as well as the most highly expressed isoform with a distal 3’ end, which we named *FENDRR2* (Fig.2C). Following transduction, RT-qPCR analysis revealed comparable overexpression for both isoforms, with an average 23-to 37-fold increase compared to empty control vector (Fig.3C). As we expected, ectopic overexpression of *FENDRR* did not significantly alter *FOXF1* expression at the RNA level, regardless of the isoform tested (Fig.3C). In contrast, we did observe a mild but significant ∼11-18% decrease in FOXF1 protein levels by western blot (Fig.3D). Together with knockdown experiments, these results indicate that *FOXF1* not only regulates *FENDRR* expression, but that *FENDRR* fine-tunes FOXF1 expression by negatively regulating its protein levels post-transcriptionally or post-translationally.

### *FENDRR* and *FOXF1* regulate a common set of genes in opposite ways

Since FOXF1 is a transcription factor and *FENDRR* appears to regulate its protein levels, we set out to gain more insights into the global transcriptome changes following knockdown of either gene. We thus performed RNA-seq following ASO-mediated knockdown of *FENDRR* and *FOXF1* and identified differentially expressed genes (DEG, *p*-adj < 0.05) compared to non-targeting ASOs. Overall, knockdown of *FENDRR* had a substantially milder transcriptomic effect than *FOXF1*, with 798 DEGs for *FENDRR* KD compared to 5,442 DEGs for *FOXF1* KD (Fig.4A,B, Supplementary Fig. S1C). To look for potential cis-regulatory effects of knocking down *FENDRR* with ASOs, we looked at changes in the expression of genes +/-1 Mb upstream and downstream of *FENDRR* TSS upon ASO-mediated knockdown compared to non-targeting ASO controls. In accordance with our RT-qPCR data, *FOXF1* mRNA levels did not significantly change upon *FENDRR* knockdown, whereas *FENDRR* was downregulated when *FOXF1* was depleted (Supplementary Fig.S1D). Only one gene, *LINC01081*, which has low expression levels, was downregulated upon *FENDRR* knockdown (Supplementary Fig.S1D, top). Notably, this same lncRNA gene was found upregulated upon *FOXF1* knockdown (Supplementary Fig.S1D, bottom).

**Figure 4.**
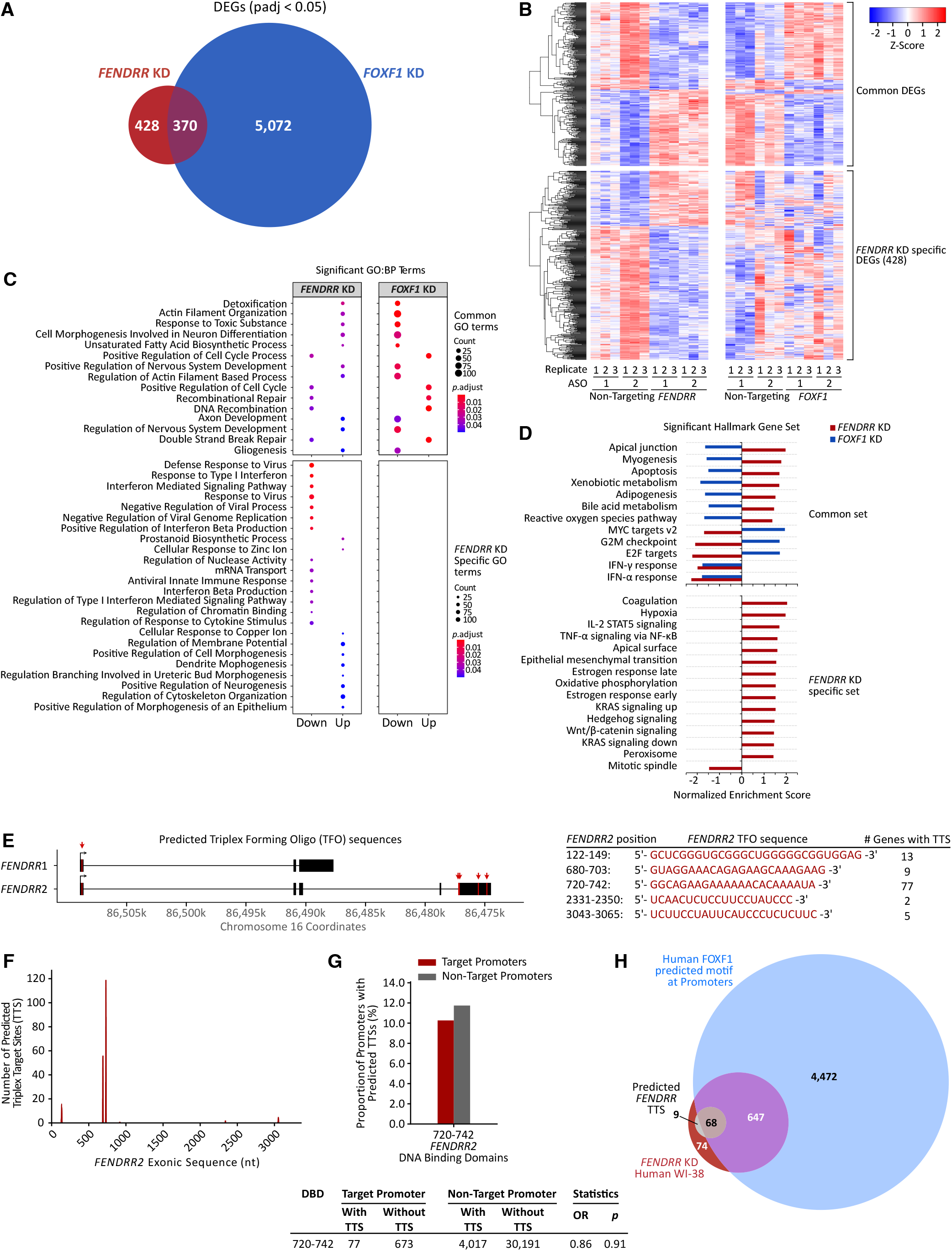
Knockdown of *FENDRR* and *FOXF1* affects a common set of genes. Differential gene expression profiling by RNA-seq. (**A**) Intersection of differentially expressed genes (DEGs; adjusted *p* <0.05) following *FENDRR* (red) and *FOXF1* (blue) depletions in WI-38 cells. (**B**) Heatmaps showing DEGs shared between *FENDRR* knockdown and *FOXF1* knockdown (top) and DEGs specific to *FENDRR* knockdown (bottom). (**C**) Gene ontology (GO) enrichment analysis (biological process) of genes upregulated (Up) or downregulated (Down) following *FENDRR* knockdown (left) and *FOXF1* knockdown (right). Top panel, overlapping GO terms; Bottom panel, terms specific to *FENDRR* knockdown. (**D**) Gene set enrichment analysis (GSEA) showing common gene sets (top) and *FENDRR-*specific gene sets (bottom). (**E**) Schematic showing the predicted positions of triplex-forming oligonucleotides (TFO) in human *FENDRR*, as analyzed by rgd-TDF (left), together with the corresponding predicted TFO sequences (right). (**F**) rgd-TDF promoter analysis predicting, for each TFO, the number of potential human *FENDRR* triplex targeting sites (TSS) at the promoters of genes differentially expressed following *FENDRR* knockdown. (**G**) The proportion of promoters from differentially expressed genes (target promoters) containing a predicted *FENDRR* TSS is lower and not significantly enriched relative to promoters of genes not differentially expressed following *FENDRR* knockdown (non-target promoters). OR, Odds ratio; TTS, Triplex Target Site; DBD, DNA Binding Domain. (**H**) Intersection of *FENDRR* KD DEGs with a predicted TTS (n=77) in their promoters with human genes containing a predicted FOXF1 binding motifs.

In accordance with the effect of *FENDRR* on FOXF1 protein levels, almost half of DEGs (46.4%; 370 DEGs) following *FENDRR* depletion were common to DEGs when *FOXF1* is knocked down (Fig.4A,B). Importantly, among this common set of DEGs, 91% were regulated in opposite manner between *FENDRR* and *FOXF1* knockdown, *i*.*e*. genes upregulated upon *FENDRR* knockdown were downregulated upon *FOXF1* depletion compared to non-targeting ASO controls and vice versa (Fig.4B). Gene ontology (GO) analysis of up- and downregulated genes upon *FOXF1* ASO-mediated depletion revealed enrichment of several expected pathways, such as embryonic morphogenesis, mesenchyme development, tissue morphogenesis, positive regulation of locomotion, and blood vessel morphogenesis (Supplementary Fig.S1E) [3, 9, 58]. When including DEGs from *FENDRR* knockdown, biological process GO-term and gene set enrichment (GSEA) analysis revealed several gene sets regulated in opposite manner to *FOXF1* depletion. Such terms included pathways associated with actin filament organization, apical junction, myogenesis, cell cycle, and neuronal development/differentiation (Fig.4C,D). When looking at GO-term and GSEA functional enrichment of genes differentially expressed specifically upon *FENDRR* knockdown (54%; 428 DEGs), terms including pathways related to antiviral responses, developmental and cellular morphogenesis, as well as key signaling cascades such as KRAS, Hedgehog, and WNT (Fig.4C,D) were enriched. While this may suggest specific pathways affected by *FENDRR* compared to FOXF1, several related terms were nonetheless found to overlap with *FOXF1* across both GO-term and GSEA analyses. For instance, while GO-term analysis indicated enrichment of antiviral response pathways upon *FENDRR* knockdown, GSEA revealed shared enrichment between *FENDRR* and *FOXF1* depletions for interferon-related signaling (Fig.4C,D). Similarly, genes involved in epithelial-to-mesenchymal transition (EMT) were found to be enriched in *FENDRR* knockdown conditions in GSEA analysis, but also for *FOXF1* DEGs in GO-term analysis (Fig.4D, Supplementary Fig.S1E). These differences likely reflect the distinct statistical frameworks underlying GSEA and GO-term analyses, as well as the possibility that different sets of genes regulated by *FENDRR* and *FOXF1* depletions converge on common pathways. Altogether, the observed transcriptomic changes between *FENDRR* and *FOXF1* ASO-mediated knockdown in embryonic lung fibroblasts support a model where *FENDRR* acts, at least in part, by negatively regulating FOXF1 protein levels.

### Most *FENDRR* knockdown DEGs have predicted FOXF1 binding motifs

Given the overlap between DEGs of *FENDRR* and *FOXF1* knockdowns, we sought to determine if any could be direct transcriptional targets of FOXF1 using a two-tiered approach. First, we reanalyzed a publicly available dataset of FOXF1 ChIP-seq from mouse E18.5 embryonic lungs [9] and intersected gene targets containing FOXF1 peaks with DEGs from both either *FENDRR* or *FOXF1* ASO-mediated knockdown, keeping only genes with one-to-one orthologs. This revealed that approximately 84% of *FOXF1* KD DEGs have FOXF1 ChIP-seq peak in the vicinity of their orthologous counterparts in mouse embryonic lung (3,380/4,047 orthologous genes; Supplementary Fig.S2A, top). Notably, strong FOXF1 peak was found at the promoter of *Fendrr* in mouse (Supplemental Fig.S2B). Together with our observation that *FENDRR* is downregulated upon *FOXF1* knockdown in WI-38 cells, these results indicate that *FENDRR* is a direct transcriptional target of FOXF1. For *FENDRR* KD DEGs, a similar proportion of gene orthologs, 84%, have a FOXF1 ChIP-seq peak near their promoter in mouse embryonic lung (529/633 genes; Supplemental Fig.S2A, bottom). Among these putative target genes, 268 genes were differentially expressed after both *FENDRR* and *FOXF1* depletions, while 261 genes were only affected after *FENDRR* knockdown.

Second, we took the FOXF1 binding motifs from JASPAR and scanned the human genome promoter sequences of DEGs from *FENDRR* and *FOXF1* knockdown to determine which may be putative targets of FOXF1. The majority (84%) of human genes with a predicted FOXF1 binding motif in their promoter overlapped with mouse orthologs that had FOXF1 ChIP-seq peaks (Supplemental Fig.S2A). Out of the 4,047 FOXF1 KD DEGs for which we could find a mouse ortholog, 2,990 genes (74%) had both a predicted FOXF1 binding motif in human and a FOXF1 ChIP-seq peak in mouse embryonic lungs at their promoter. An additional 575 had only a human predicted FOXF1 binding motif and 390 had only a FOXF1 ChIP-seq peak in its mouse ortholog. Importantly, from the 633 *FENDRR* KD DEGs for which we could find a mouse ortholog, 476 genes (75%) had both a predicted FOXF1 binding motif in human and a FOXF1 ChIP-seq peak in mouse embryonic lungs at their promoter. Another 89 genes had only a human predicted FOXF1 binding motif and 53 had only a FOXF1 ChIP-seq peak in its mouse ortholog. Together, these data suggest that a significant subset of differently expressed genes upon *FENDRR* loss may be directly regulated by FOXF1.

### Human *FENDRR* lacks evidence of triplex-forming potential

In mouse, *Fendrr* is mainly localized in the nucleus and associated with the chromatin fraction. Previous studies have proposed that mouse *Fendrr* transcripts can form RNA-DNA triplexes at the promoters of a subset of genes to regulate their expression (20 out of 60 DEGs in *Fendrr* knockout lungs) [20, 46]. Since human *FENDRR* is localized both in the cytoplasm and nucleus (Fig.2E,F), it could also function to regulate genes by forming RNA-DNA triplexes. However, pairwise sequence alignments of human and mouse *FENDRR* transcripts reveal that predicted mouse triplex forming oligonucleotide (TFO) sequences is poorly conserved in human (Supplemental Files 1,2). Nevertheless, it remains possible that other sequences in human *FENDRR* transcripts act as triplex forming oligos. To identify such sequences, we used the *Triplex Domain Finder* (TDF) algorithm [39, 40] to characterize the RNA-DNA triplex-forming potential of *FENDRR1* and *FENDRR2* isoforms and test whether promoters of differentially expressed genes after *FENDRR* knockdown are enriched in triplex targeting sites (TTS). This analysis predicted only 1 TFO in the 5’ end of *FENDRR1*, whereas 4 additional TFOs were identified in *FENDRR2*, only one of which (nt 719-742) was significant (Fig.4E-G). For this sequence, the TDF algorithm detected triplex target sites (TTS) in the promoter of only 10.3% (77/750) of genes differentially expressed in *FENDRR* depleted WI-38 cells (Fig.4F,G). This proportion was not statistically significant compared to TTS found in the promoter of other genes of the genome (Fig.4G), *i*.*e*. expressed genes not differentially regulated when *FENDRR* is depleted. Notably, no TTS were detected in the FOXF1 promoter, supporting our observation that human *FENDRR* does not regulate *FOXF1* transcription. In addition, 42/77 (55%) genes with a predicted *FENDRR* TTS in their promoter are differentially expressed under both *FENDRR* and *FOXF1* knockdown conditions, 57/77 (74%) have a *FOXF1* ChIP-seq peak in their promoter in mouse lungs, and 68/77 (88%) have a predicted human FOXF1 motif in their promoter (Fig.4H), suggesting that changes in their expression following *FENDRR* ASO-mediated knockdown are likely due to changes in FOXF1 protein levels. Together, these results suggest that, in contrast to what was reported in mice, human *FENDRR* does not appear to have triplex forming potential at promoters of differentially expressed genes.

### *FENDRR* and *FOXF1* regulate fibroblast-to-myofibroblast transition in opposite ways

To further understand the interplay between *FENDRR* and *FOXF1*, we evaluated the cellular impact of knocking down each gene on human embryonic lung fibroblasts. FOXF1 has been shown to regulate fibroblast migration and fibroblast-to-myofibroblast transition [5, 6, 58]. *FENDRR* was recently shown to play a role in idiopathic pulmonary fibrosis and asbestos-induced fibrotic lesions [21, 22]. Thus, we first conducted wound healing assays to determine the role of *FENDRR* and FOXF1 on fibroblast migration. While ASO-mediated knockdown of FOXF1 led to reduced migratory capacity compared to non-targeting ASO controls, as expected (*REF*), knockdown of *FENDRR* did not significantly alter migration (Fig.5A). Accordingly, lentivirus-mediated overexpression of either *FENDRR1* or *FENDRR2* isoforms did not affect cell migration either (Fig.5B).

**Figure 5.**
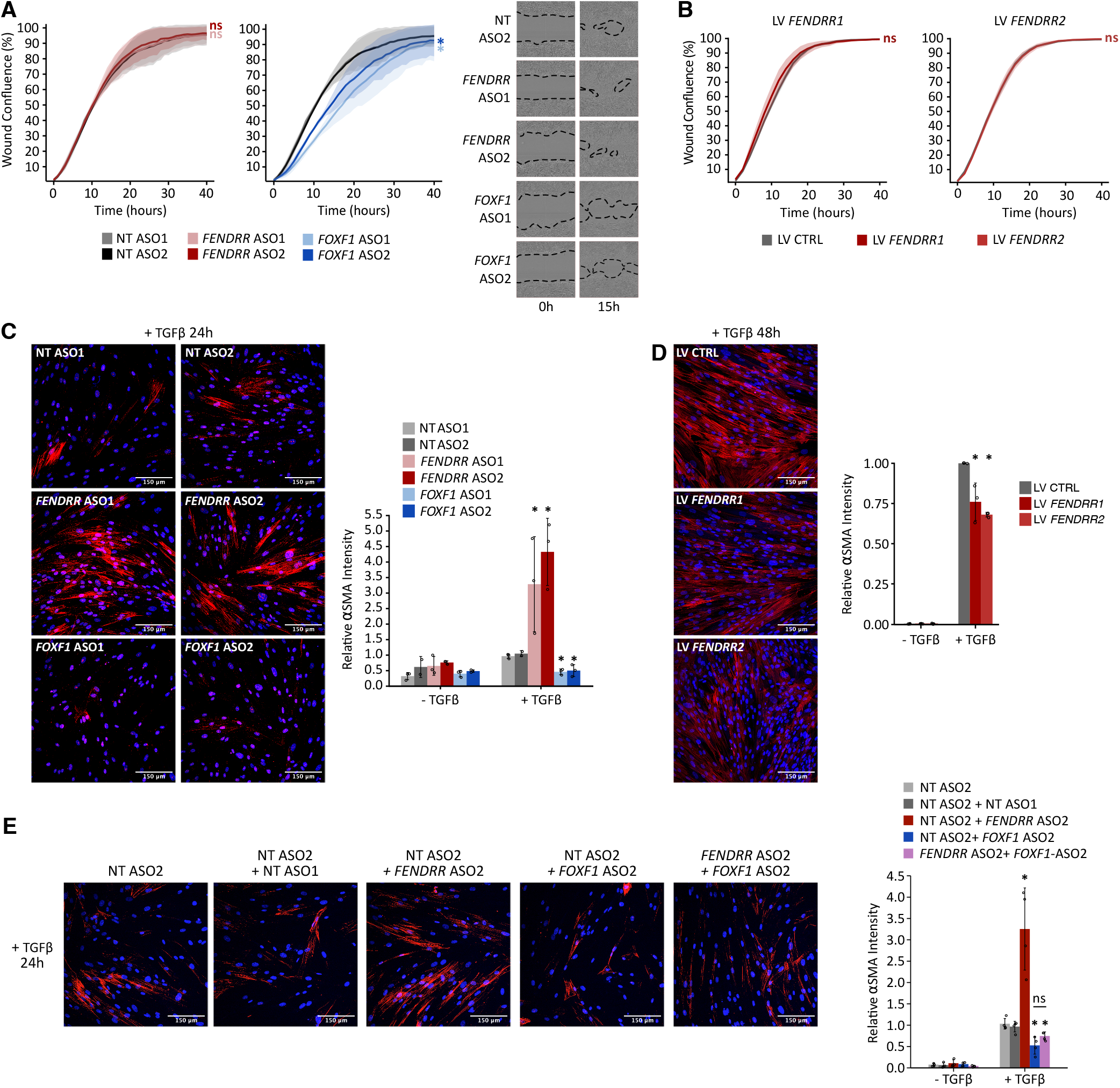
*FENDRR* and *FOXF1* exert opposing effects on lung fibroblast-to-myofibroblast transition. (**A**) Quantification of scratch wound healing following *FENDRR* (left; red) and *FOXF1* (middle; blue) knockdown in WI-38 cells. Wound closure was monitored by real-time quantitative live-cell imaging. Representative images at 0h and 15h are shown on the right. (**B**) Quantification of scratch wound healing in WI-38 overexpressing *FENDRR* isoforms via lentiviral transduction, compared with empty-vector control cells. Wound closure was monitored by real-time quantitative live-cell imaging. For both (A) and (B) data are presented as the percentage of wound area occupied by cells, normalized to 0h. Lines represent the mean of three biological replicates, and error bars indicate standard deviation (SD). Statistical significance was determined by two-way ANOVA followed by Dunnett’s post hoc test (\**p-*val < 0.05; ns, not significant). (**C**) Representative immunofluorescence staining of α-smooth muscle actin (αSMA), with signal quantification across three biological replicates, to assess fibroblast-to-myofibroblast transition (FMT) 24h after TGFβ1 treatment and ASO-mediated knockdown of *FENDRR* or *FOXF1* in WI-38 cells. Signal intensities were calculated relative to TGFβ1-treated control antisense oligonucleotides (NT ASO1 and NT ASO2). Bars represent the mean, dots indicate individual replicates, and error bars denote standard deviation (SD). Statistical significance was determined by one-way ANOVA followed by Dunnett’s post hoc test (\**p*-val < 0.05; ns, not significant). (**D**) Representative immunofluorescence staining of αSMA with signal quantification across three biological replicates, to assess FMT 48h after TGFβ1 treatment in WI-38 cells overexpressing *FENDRR* isoforms via lentiviral transduction. Signal intensities were calculated relative to TGFβ1-treated empty-vector control (LV CTRL). Bars represent the mean, dots indicate individual replicates, and error bars show standard deviation (SD). Statistical analysis was performed as in C. (**E**) Immunofluorescence staining of alpha smooth muscle actin (αSMA) marker to assess FMT in WI-38 cells treated (right) or not (left) with TGFß1 for 48h and following ASO-mediated knockdown of *FENDRR, FOXF1*, or both, compared to non-targeting ASO controls (NT) across four biological replicates. Quantification of αSMA Intensity relative to non-targeting control (NT ASO2) is shown on the right. DAPI was used to stain nuclei.

RNA sequencing results revealed several gene sets involved in myogenesis, epithelial-to-mesenchymal transition, and cell architecture regulated in opposite ways between *FENDRR* and *FOXF1* depleted WI-38 cells (Fig.4C,D). We also showed that both *FENDRR* and *FOXF1* RNA levels are downregulated by TGFβ1 (Fig.1F), one of the primary factors driving differentiation of fibroblasts into contractile myofibroblasts [59]. Therefore, we treated *FENDRR* and *FOXF1* depleted WI-38 cells with TGFβ1 to induce fibroblast-to-myofibroblast transition (FMT) and measured α-smooth muscle actin (αSMA) stress fiber formation, a hallmark of FMT [60], by immunofluorescence. FMT induction was monitored over time, revealing partial stress fiber formation at 24h and much more advanced formation by 48h (Supplementary Fig.S3A). Treatment of WI-38 cells with 10 ng/ml TGFβ1 for 24 h following ASO-mediated knockdown of *FENDRR* and *FOXF1* revealed opposite effects on FMT. *FENDRR* depletion showed increased αSMA stress fiber formation compared to non-targeting ASO controls, while *FOXF1* knockdown abrogated FMT (Fig.5C, Supplemental Fig.S3B). Moreover, simultaneous knockdown of *FOXF1* and *FENDRR* in WI-38 cells abolished αSMA stress fiber formation, indicating that the increased FMT following *FENDRR* depletion is dependent on *FOXF1* (Fig.5E, Supplemental Fig.S4A). TGFβ1 treatment was required to induce fibroblast-to-myofibroblast, as knockdown of *FENDRR* alone did not lead to increased αSMA stress fiber formation (Supplementary Fig.S3B, S4A). This suggests a role for *FENDRR* in myofibroblast activation and the priming of differentiation programs rather than directly driving myofibrogenesis. This is supported by the upregulation of key genes associated to fibroblast activation and fibrogenesis upon *FENDRR* knockdown in the absence of TGFβ1 (Supplementary Fig.S3C), while changes in the expression of canonical markers of differentiated myofibroblasts, including *ACTA2, FN1, COL1A1*, and *COL1A2*, remained not significant.

Given that *FENDRR* overexpression lowers FOXF1 protein levels, a corresponding decrease in FMT would be expected. Accordingly, we found that lentivirus-mediated overexpression of *FENDRR1 and FENDRR2* isoforms led to a significant decrease in αSMA stress fiber formation 48h after TGFβ1 treatment (Fig.5D, Supplemental Fig.S4B). Taken together, these data indicate that *FENDRR* and *FOXF1* have opposing effects, where *FOXF1* acts as a positive regulator of FMT, whereas *FENDRR* counteracts this process, likely by downregulating FOXF1 protein level.

## DISCUSSION

Proper lung development relies on tightly coordinated gene regulatory programs within the mesenchyme, where variations in transcription factor dosage can result in profound developmental consequences [7, 11, 61, 62]. FOXF1 is a central component of these programs, with genetic and experimental evidence demonstrating that both reduced and excessive FOXF1 activity perturb lung morphogenesis [7, 9, 11], and that heterozygous mutations or regulatory deletions cause alveolar capillary dysplasia with misalignment of pulmonary veins (ACDMPV) [10–12]. In this study, we examined the relationship between FOXF1 and the divergently transcribed long noncoding RNA *FENDRR* in embryonic lung fibroblasts. Our results support a model in which FOXF1 transcriptionally activates *FENDRR*, while *FENDRR* feeds back to dampen FOXF1 at the protein level, thereby establishing a rheostat-like mechanism that influences embryonic lung fibroblasts transcriptional programs and behavior. While our findings do not establish a fully defined molecular mechanism, they point to an additional layer of regulation that may contribute to the dosage sensitivity of FOXF1 during lung development.

Single-cell transcriptomic analyses confirm that *FENDRR* and *FOXF1* are co-expressed across overlapping mesenchymal and mesodermal cell types in human and mouse embryonic lungs, including fibroblasts, smooth muscle cells, endothelial cells, and mesothelial cells. Within fibroblast populations, both genes display gradients of expression that are higher in proximal regions and decrease distally, consistent with spatial patterning of mesenchymal signaling pathways during lung development [28]. Induction of Sonic Hedgehog and TGFβ signaling in embryonic lung fibroblasts further indicates that both genes respond similarly to upstream developmental cues, supporting the view that they are transcriptionally co-regulated at least under certain conditions.

Despite this conserved co-expression pattern, our data reveal notable differences between human and mouse *FENDRR* at the level of isoform diversity and subcellular localization. Human *FENDRR* exhibits multiple alternatively processed isoforms and is detected in chromatin-associated and cytoplasmic compartments, whereas mouse *Fendrr* is predominantly nuclear and chromatin-enriched. These differences are consistent with earlier observations that lncRNAs can retain positional conservation while diverging substantially in primary sequence, processing, and subcellular localization [63]. Accordingly, our results suggest that conclusions drawn from murine *Fendrr* models may not be directly generalizable to human *FENDRR*, particularly with respect to molecular mechanism.

A principal observation of this study is that perturbation of *FENDRR* levels affects FOXF1 protein abundance while leaving *FOXF1* mRNA largely unchanged. Specifically, *FENDRR* depletion leads to a reproducible ∼30% increase in FOXF1 protein, whereas ectopic expression of distinct *FENDRR* isoforms produces the opposite effect. These reciprocal effects, although not of high magnitude, are consistent across knockdown and overexpression approaches and across multiple *FENDRR* isoforms. Our results are also consistent with genetic observations from mouse *Fendrr* knockouts lungs, where *Foxf1* mRNA levels remain largely unchanged [19, 20]. By contrast, *FOXF1* depletion reduces *FENDRR* RNA levels, and ChIP-seq indicate that FOXF1 binds the *FENDRR* promoter in mouse lung and human gastrointestinal tumors [9, 64], supporting the notion that *FENDRR* is a transcriptional target of FOXF1. This is further supported by previous studies showing a decrease in *FENDRR* expression in ACDMPV patient lung with heterozygous point mutation of *FOXF1* [57].

Together, these findings are compatible with a regulatory relationship in which FOXF1 promotes *FENDRR* transcription, while *FENDRR* feeds back to constrain FOXF1 protein abundance. While many lncRNAs have been shown to regulate the transcription of neighboring protein coding gene in *cis* [65], examples where the lncRNA regulates the protein-level of its neighboring gene have been few. Such protein-level modulation is particularly relevant for dosage-sensitive transcription factors. Recent studies demonstrate that transcription factor concentration–response relationships are often highly nonlinear, such that modest changes in protein levels can preferentially affect subsets of target genes [66–69]. By buffering FOXF1 protein abundance, *FENDRR* may therefore stabilize mesenchymal gene expression programs during narrow developmental windows, preventing inappropriate activation or repression of FOXF1-responsive loci. However, the underlying mechanism by which *FENDRR* affects FOXF1 protein levels remains unresolved. While our data indicate the directionality of the regulatory interaction, it does not distinguish between effects on translation, protein stability, or indirect regulation through intermediate factors.

Comparative transcriptomic analyses reveal that *FENDRR* knockdown produces a more limited set of differentially expressed genes than *FOXF1* knockdown, consistent with FOXF1 functioning as a transcription factor and *FENDRR* acting indirectly. Nearly half of the genes altered upon *FENDRR* depletion overlap with FOXF1-dependent genes, and the majority of these shared targets are regulated in opposite directions between the two conditions. This pattern is consistent with the observed inverse effects of *FENDRR* on FOXF1 protein levels and suggests that at least a subset of *FENDRR*-dependent transcriptional changes may reflect secondary consequences of altered FOXF1 activity. Supporting this interpretation, many genes responsive to *FENDRR* depletion contain predicted FOXF1 binding motifs and/or overlap with FOXF1 ChIP-seq peaks identified in orthologous genes in mouse embryonic lung tissue. At the same time, *FENDRR* depletion also affects genes not detectably associated with FOXF1 binding, indicating that additional FOXF1-independent effects cannot be excluded. Thus, while the data are consistent with a model in which *FENDRR* modulates FOXF1-dependent transcriptional output, they do not imply that all functions of *FENDRR* are mediated exclusively through FOXF1.

Previous studies in mouse have proposed that *Fendrr* can modulate gene expression through RNA– DNA triplex formation at target promoters [20, 46]. In contrast, our analyses find limited evidence that human *FENDRR* exhibits triplex-forming potential at promoters of genes affected by *FENDRR* knockdown. Predicted triplex-forming oligo sequences are sparse, poorly conserved, and not significantly enriched among *FENDRR*-dependent genes. While these results argue against widespread triplex-mediated regulation by human *FENDRR* in this cellular context, they do not exclude more restricted or condition-specific roles for DNA-associated interactions. Nevertheless, the data support the idea that human *FENDRR* may act predominantly through mechanisms distinct from those described in mouse.

Functionally, we observe opposing effects of *FENDRR* and *FOXF1* on TGFβ-induced fibroblast-to-myofibroblast transition (FMT). *FOXF1* depletion markedly impairs αSMA stress fiber formation, consistent with previous reports that FOXF1 is required for mesenchymal differentiation responses. In contrast, *FENDRR* depletion enhances FMT, while *FENDRR* overexpression attenuates it. Importantly, the increased FMT observed upon *FENDRR* loss is abolished when *FOXF1* is simultaneously depleted, indicating that this effect depends on FOXF1 activity.

These findings have clear implications for fibrotic lung disease. FOXF1 is upregulated in fibroblasts from idiopathic pulmonary fibrosis (IPF) patients [23], whereas *FENDRR* expression is reduced, and ectopic *FENDRR* expression attenuates fibrotic remodeling in mouse lungs [21, 22]. Notably, we show that *FENDRR* depletion alone is insufficient to drive myofibroblast differentiation in the absence of TGFβ signaling. Thus, our data provide a mechanistic framework linking these observations, suggesting that loss of *FENDRR*-mediated buffering may shapes the threshold or extent of FOXF1-dependent responses, thereby influencing the sensitivity of fibroblasts to profibrotic cues such as TGFβ rather than acting as a primary determinant of differentiation. This could also have implications for ACDMPV where deletions involving the *FENDRR* locus could affect levels of FOXF1, thereby contributing to differences in pathological manifestations.

Several limitations of this study warrant consideration. First, most functional experiments were conducted in the WI-38 human embryonic lung fibroblast line, which, while developmentally relevant, may not fully capture the cellular heterogeneity and spatial context of the embryonic lung *in vivo*. Second, while changes in FOXF1 protein abundance are reproducible, the precise molecular mechanism of *FENDRR*, whether through modulation of translation, protein stability, or interaction with specific cofactors, remains unresolved. Third, while we document isoform diversity of human *FENDRR*, our functional analyses do not exhaustively dissect isoform-specific roles beyond two representative transcripts.

Future work will be required to delineate how *FENDRR* influences FOXF1 protein abundance and whether this interaction occurs directly or through intermediary pathways. It will also be important to assess whether similar regulatory relationships operate in other *FOXF1*-expressing mesenchymal populations and during later stages of lung development or disease. More broadly, these findings contribute to accumulating evidence that lncRNAs can participate in fine-scale quantitative regulation of transcription factor activity, adding complexity to gene dosage control during development.

In summary, our data support a model in which *FENDRR* and FOXF1 participate in a rheostat-like system that modulates FOXF1 protein levels and influences transcriptional responses in embryonic lung fibroblasts. While the effects on protein abundance are not of high magnitude, they may nonetheless be relevant in developmental contexts where FOXF1 dosage must be tightly constrained with implications for lung fibrosis, and ACDMPV pathogenesis.

## ACKNOWLEDGEMENTS

The authors thank Sarah Boissel, Julie Lord, Dominic Filion and their team at the IRCM Functional Genomics, Flow Cytometry, Microscopy core facilities for their valuable technical support. Tha authors also thank the Université de Sherbrooke RNomics platform for the Nanopore direct RNA library preparation and sequencing as well as Daniel Zenklusen and Pascal Raymond (Université de Montréal) for their expertise on smFISH imaging.

## SUPPLEMENTARY DATA

Supplementary data are available Online.

## CONFLICT OF INTEREST

The authors declare no competing interests.

## FUNDING

This work was supported by a Fonds de Recherche du Québec - Santé (FRQS) New Investigator Starting Grant (265520), a Canadian Institutes of Health Research (CIHR) Project grant (401286), IRCM Foundation startup funds, as well as a Canadian Foundation for Innovation (CFI) John-R. Evans Leaders fund (37709) to M.S. J.F.L. is supported by a Fonds de Recherche du Québec - Nature & Technologies (FQRNT) PhD scholarship (355015), a Simon-Pierre Noël Award from Université de Montréal, and an Angelo-Pizzagalli award from the IRCM Foundation. M.S. is a recipient of Fonds de Recherche du Québec - Santé (FRQS) Research Scholar Junior 1 and Junior 2 awards (265520, 330732).

## DATA AVAILABILITY

The short-read RNA and long-read Nanopore direct RNA sequencing data are deposited in GEO. Single-cell RNA-seq data from human and mouse embryonic lungs were obtained from ArrayExpress under accession E-MTAB-11278 and GEO under accession GSE149563, respectively. The mouse FOXF1 ChIP-seq data were retrieved from GEO under accession GSE77951.

## AUTHOR CONTRIBUTIONS STATEMENTS

Conceptualization: MS and JFL; Methodology: MS, JFL, SM, and SE; Investigation: JFL, SM, SE, EG, and MS; Computational Analyses: JFL, GDB, SM, SE, and VC; Experimental Analyses: JFL, MS, SM, SE, EG, GDB, and MM; Writing Draft: JFL and MS; Figures: JFL, GDB, and MS; Supervision, Administration, Funding Acquisition, and Resources: MS; All authors have read and approved the final manuscript.

## Notes

### Competing Interest Statement

The authors have declared no competing interest.

