## Supplemental Figures for "The lncRNA *FENDRR* fine-tunes FOXF1 protein levels through a negative feedback loop governing human embryonic lung fibroblast-to-myofibroblast transition"

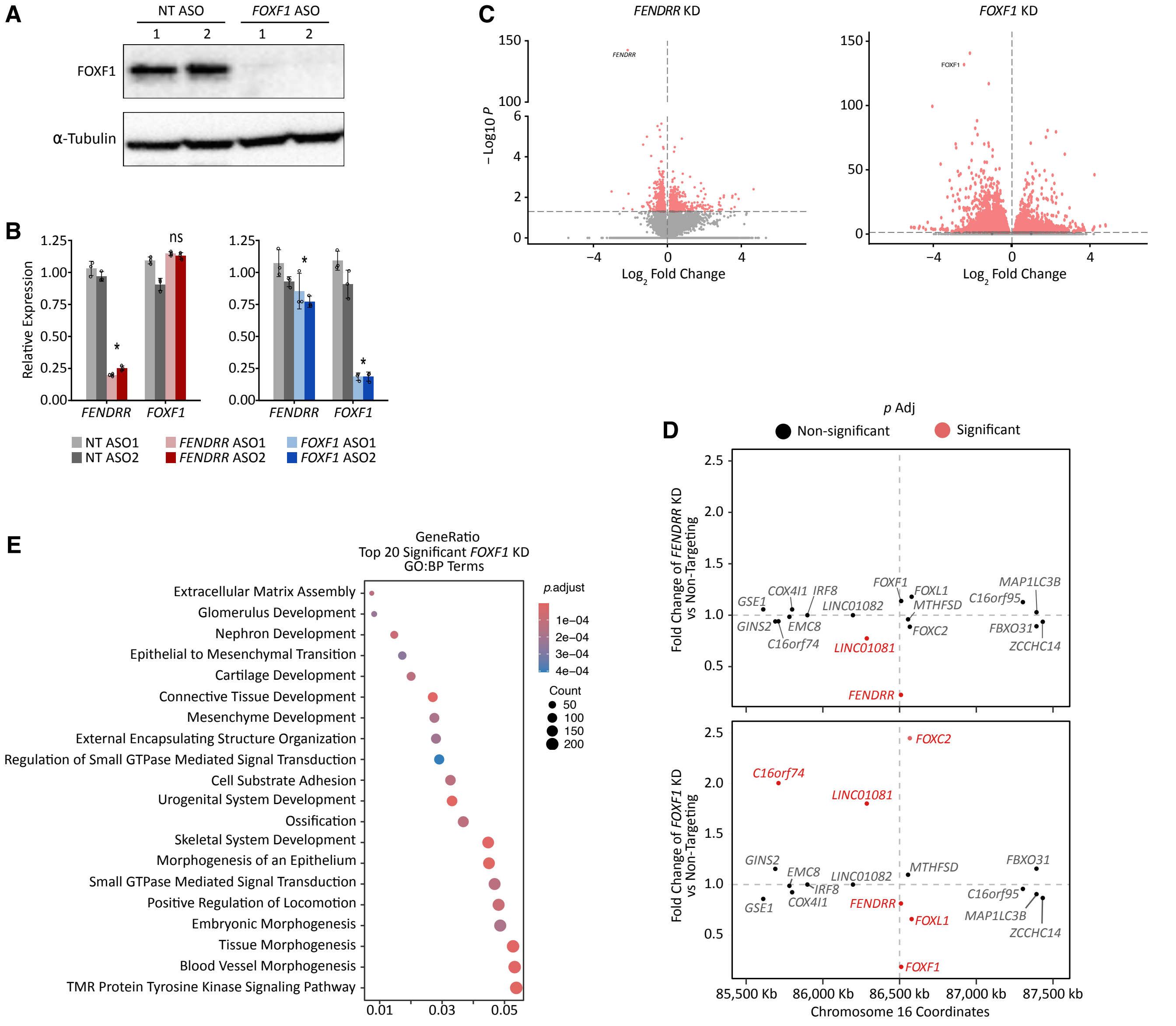


**Supplemental Figure S1. Effect of *FENDRR* and *FOXF1* depletion on WI-38 transcriptome.** (**A**) Representative westerm blot of FOXF1 and α-Tubulin protein levels 48h after ASO-mediated *FOXF1* knockdown in WI-38. (**B**) *FENDRR* and *FOXF1* RNA levels assessed by RNA-seq 48h after ASO-mediated *FENDRR* or *FOXF1* knockdown in WI-38. (**C**) Volcano plots of whole transcriptomic changes 48h after *FENDRR* (left) and *FOXF1* (right) ASO-mediated knockdown in WI-38. (**D**) Expression levels from RNA-seq of genes within 1 Mb upstream and downstream of *FENDRR*'s transcription start site (TSS) 48h after *FENDRR* (top) and *FOXF1* (bottom) ASO-mediated knockdown. (**E**) Top 20 significant GO terms (biological process) following *FOXF1* depletion. Abbreviation: NT, Non-targeting; ns, not significant; FC, fold change.

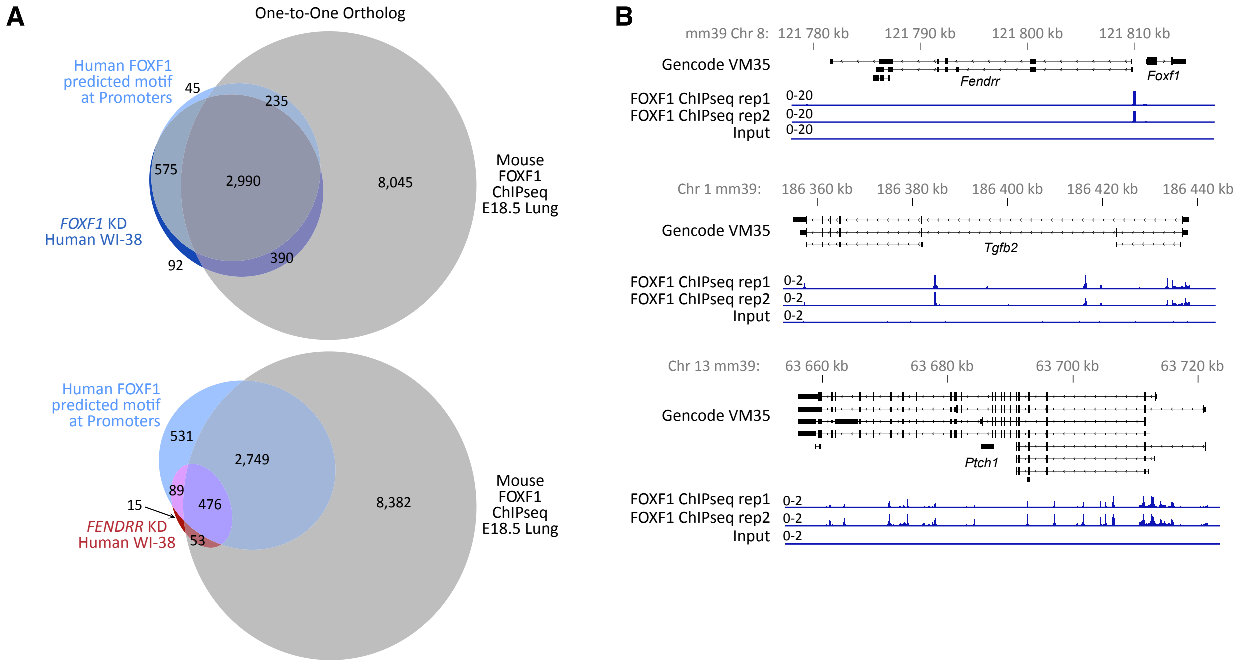


**Supplementary Figure S2. Overlap between differentially expressed genes and FOXF1 ChIP-seq.** FOXF1 ChIP-seq peaks from mouse E18.5 embryonic lungs were analyzed to identify target genes based on each peak's nearest feature (gene). Differentially expressed genes (DEG) from FENDRR and FOXF1 ASO-mediated knockdown in human WI-38 embryonic lung fibroblasts were intersected with mouse ortholog genes associated with FOXF1 ChIP-seq peaks. (**A**) Euler diagrams representing the intersect between human WI-38 *FOXF1* knockdown (top) or *FENDRR* knockdown (bottom) DEGs, human genes with predicted FOXF1 binding motifs in their promoter, and mouse ortholog genes (one-to-one) associated with FOXF1 ChIP-seq in E18.5 embryonic lungs. (**B**) Representative examples of FOXF1 ChIP-seq peaks identified in mouse E18.5 embryonic lung near mouse orthologs of differentially expressed genes (*Fendrr*, *Tgfb2*, and *Ptch1*) following *FENDRR* and *FOXF1* ASO-mediated knockdown in human WI-38 embryonic lung fibroblasts. Genomic coordinates (mm39) and Gencode vm35 annotations were used.

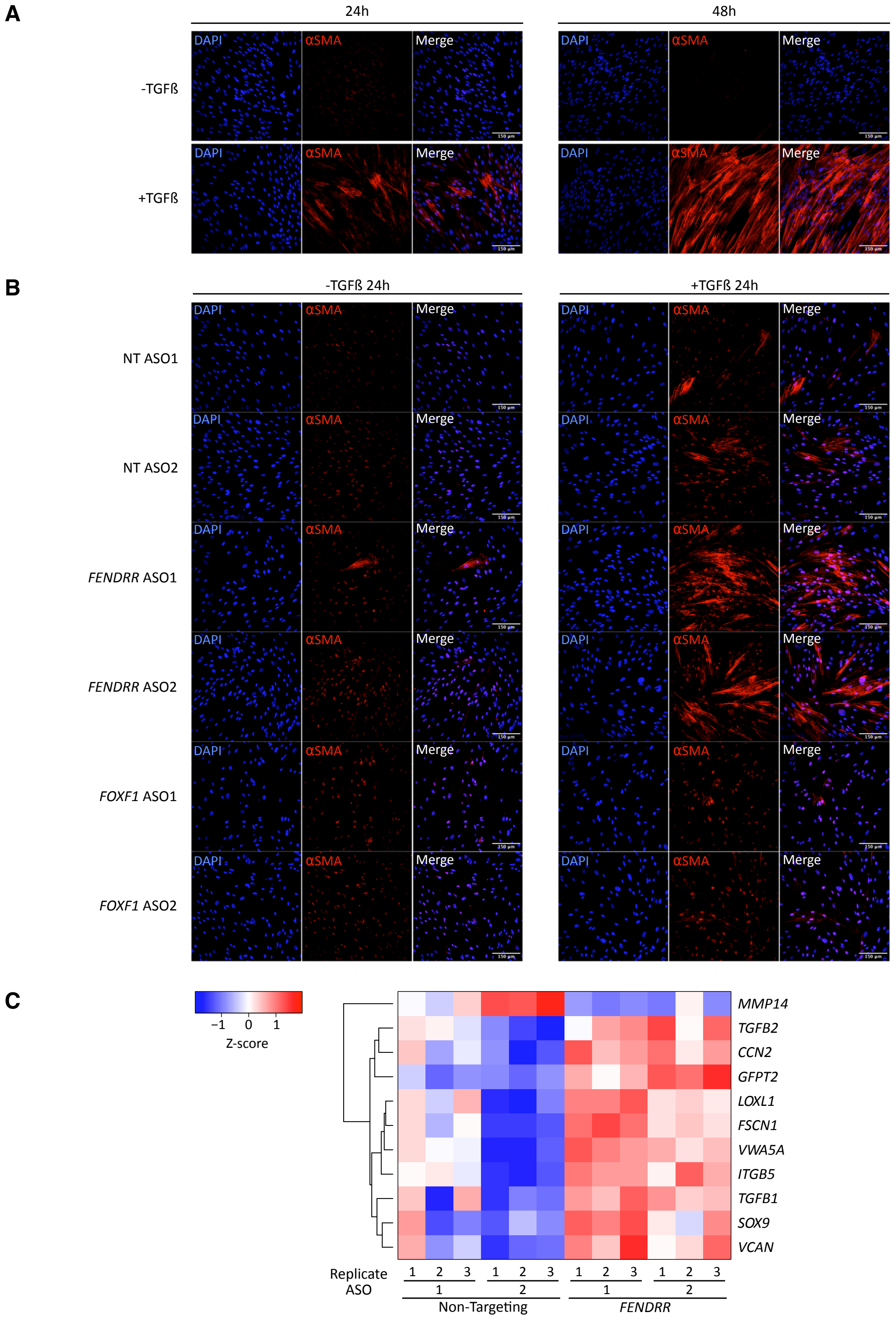


**Supplementary Figure S3. Fibroblast-to-myofibroblast transition following *FENDRR* or *FOXF1* knockdown.** (**A**) Immunofluorescence staining of alpha smooth muscle actin (αSMA) marker to assess fibroblast-to-myofibroblast transition (FMT) following 24h (left) or 48h (right) TGFß1 treatment (10 ng/ml) in WI-38. (**B**) Immunofluorescence staining of αSMA to assess FMT of WI-38 treated with 10 ng/ml TGFß1 for 24h (right) or not (left) after ASO-mediated knockdown of *FENDRR* or *FOXF1*. (**C**) Heatmap from RNA-seq data of differentially expressed genes associated with TGFß1-mediated fibroblast activation and fibrogenesis following ASO-mediated knockdown of *FENDRR* in WI-38 cells.

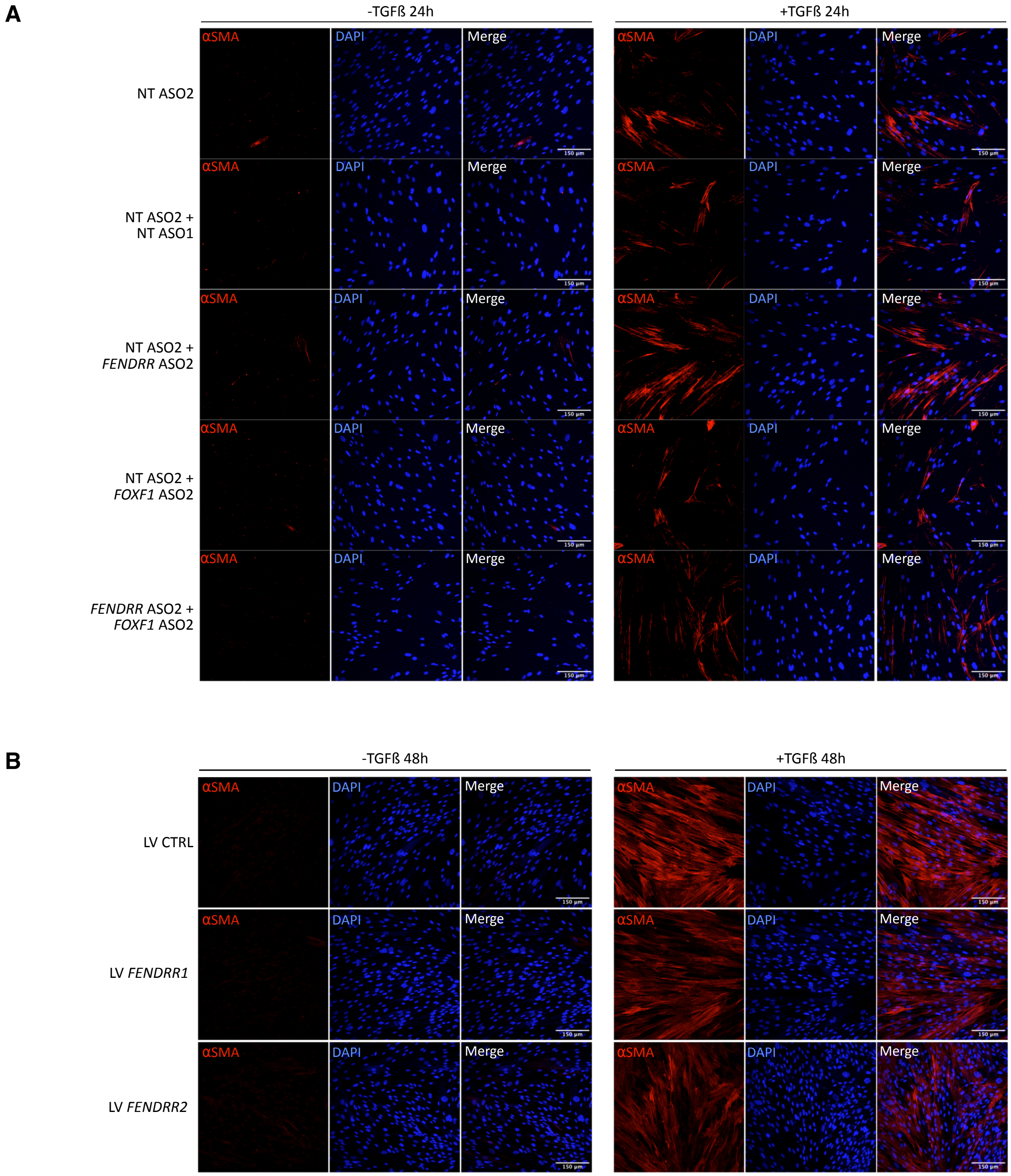


**Supplementary Figure S4. Fibroblast-to-myofibroblast transition following *FENDRR* and *FOXF1* double knockdown and *FENDRR* overexpression.** (**A**) Immunofluorescence staining of alpha smooth muscle actin (αSMA) marker to assess fibroblast-to-myofibroblast (FMT) in WI-38 cells treated (right) or not (left) with TGFß1 for 48h and following ASO-mediated knockdown of *FENDRR, FOXF1*, or both, compared to non-targeting ASO controls (NT). (**B**) Immunofluorescence staining of αSMA stress fiber marker to assess fibroblast-to-myofibroblast (FMT) in WI-38 cells treated (right) or not (left) with TGFß1 for 48h and following lentiviral-mediated overexpression of *FENDRR* isoforms compared to empty lentivirus control (LV CTRL). DAPI was used to stain nuclei.
